# Chromosome end protection by RAP1-mediated inhibition of DNA-PK

**DOI:** 10.1101/2024.12.28.630583

**Authors:** Patrik Eickhoff, Ceylan Sonmez, Charlotte E.L. Fisher, Oviya Inian, Theodoros I. Roumeliotis, Angela dello Stritto, Jörg Mansfeld, Jyoti S. Choudhary, Sebastian Guettler, Francisca Lottersberger, Max E. Douglas

## Abstract

During classical non-homologous end joining (cNHEJ), DNA-dependent protein kinase (DNA-PK) encapsulates free DNA ends, forming a recruitment platform for downstream end-joining factors including Ligase 4 (LIG4)^1^. DNA-PK can also bind telomeres and regulate their resection^2–4^, but does not initiate cNHEJ at this position. How the end joining process is regulated in this context-specific manner is currently unclear. Here we show that the shelterin components TRF2 and RAP1 form a complex with DNA-PK that directly represses its end joining function at telomeres. Biochemical experiments and cryo-electron microscopy reveal that when bound to TRF2, RAP1 establishes a network of interactions with KU and DNA that prevents DNA-PK from recruiting LIG4. In mouse and human cells, RAP1 is redundant with the Apollo nuclease in repressing cNHEJ at chromosome ends, demonstrating that the inhibition of DNA-PK prevents telomere fusions in parallel with overhang-dependent mechanisms. Our experiments show that the end joining function of DNA-PK is directly and specifically repressed at telomeres, establishing a molecular mechanism for how individual linear chromosomes are maintained in mammalian cells.

---

Mammalian telomeres are protected from cNHEJ by the shelterin subunit TRF2, which is proposed to hide the chromosome end from DNA repair factors by forming a lariat structure referred to as a t-loop^2,5,6^. T-loop assembly requires a terminal 3’ overhang^7^, which at blunt telomeres resulting from leading strand DNA replication, is formed via a 5’ resection step mediated by Apollo exonuclease^8–12^. Surprisingly, DNA-PK is required for this resection step, but is unable to activate cNHEJ even after Apollo deletion^13^, suggesting a mechanism is in place to directly block the end joining process at telomeres. We hypothesised this mechanism may involve the conserved shelterin subunit RAP1. In budding yeast, Rap1 binds directly to telomeres and protects them from cNHEJ^14^. However, in mammalian cells, RAP1 is recruited via TRF2 and its role in end protection remains elusive^15–19^.

To examine this idea, we employed telomere-fluorescence in situ hybridisation (FISH) to measure chromosome fusions in conditional Apollo knockout (KO) mouse embryonic fibroblasts (MEFs) with or without RAP1 (Fig. S1a). In agreement with previous studies, Apollo-deleted MEFs showed some chromatid fusions that were LIG4-independent and so not caused by cNHEJ (Figs. 1a-c, compare –sgRap1, ±Cre)^13^. No fusions were observed upon Crispr-mediated deletion of RAP1 in Apollo proficient MEFs, also consistent with previous work^19–21^ (Figs. 1a-c, compare ±sgRap1, –Cre). However, when RAP1 and Apollo were deleted together, approximately 15 % of telomeres per metaphase were engaged in LIG4-dependent chromosome-type fusions (Figs. 1a and c, +sgRap1, +Cre). A similar effect was induced by mutating the binding sites for Apollo and RAP1 on TRF2, and the resulting fusions were prevented by deleting the iDDR (Fig. S1b-d), which restores telomeric overhangs in the absence of Apollo^13^. The data above strongly suggests that 3’ overhangs and RAP1 can each protect mouse telomeres from cNHEJ. To test whether this is also the case for human telomeres, we repeated the analysis in non-transformed human cells. Unlike in cancer cell lines^22,23^ telomere fusions were not induced in p53-/-RPE-1 cells lacking only Apollo (Figs. 1d, e and S1g). However, consistent with the data above, 20-30 % of telomeres fused in a DNA-PK-dependent manner when RAP1 was also removed (Figs. 1d, e, Fig. S1e-h). A proportion of these fusions involved only one telomere per chromosome end and in line with the established role of Apollo, these telomeres had exclusively been replicated as the leading strand (Fig. S1g and i). We conclude that in mouse and human cells, telomeres are protected from cNHEJ by two equally effective and parallel pathways: 3’ overhangs and the presence of RAP1.

**Figure 1.**
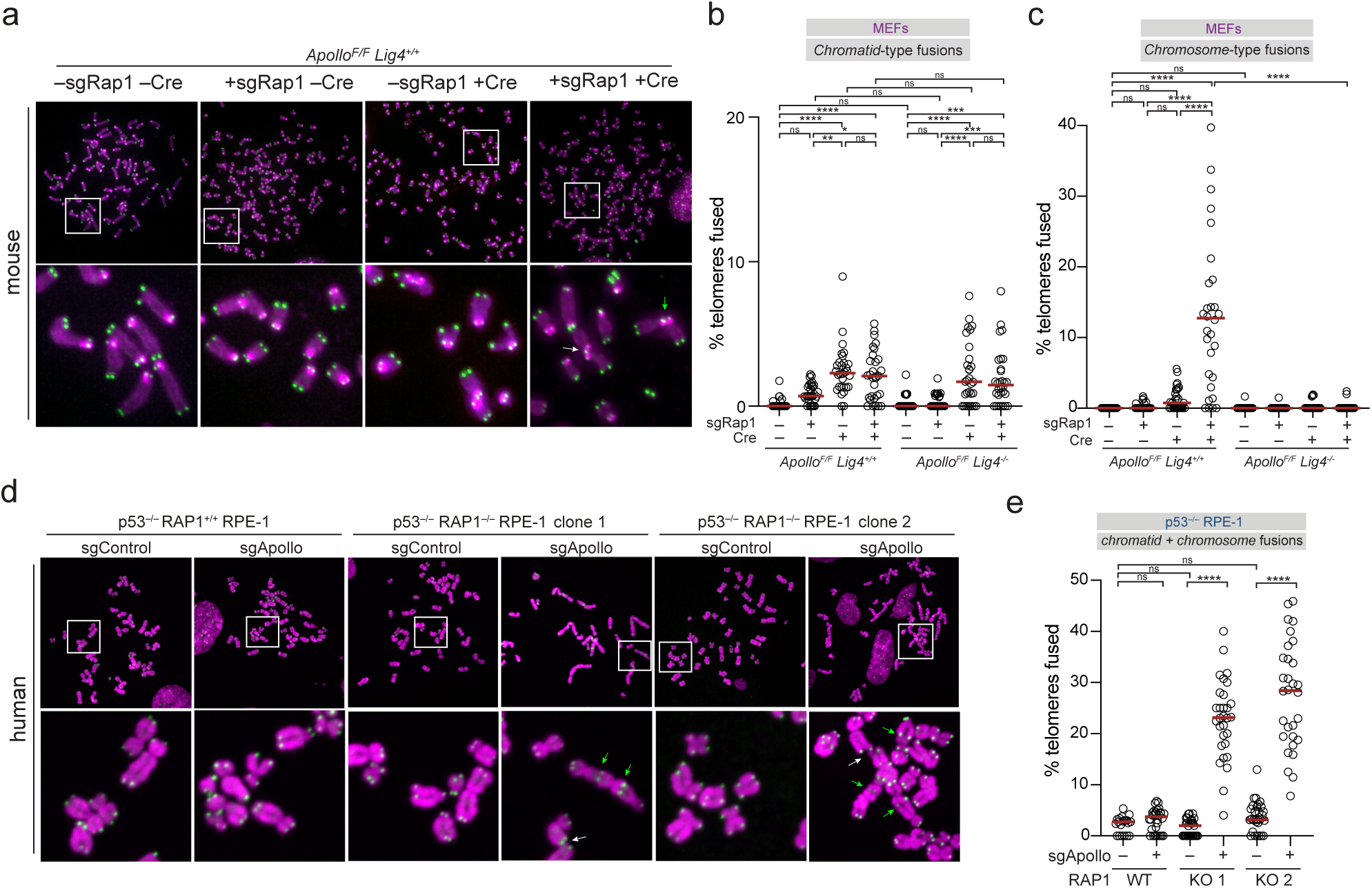
RAP1 and APOLLO redundantly prevent cNHEJ at telomeres in mouse and human cells. **a.** Represent-ative FISH of metaphase spreads of ApolloF/F Lig4+/+ MEFs 108 h after transduction with sgRap1 and/or Hit & Run Cre. Telomeres were detected with Cy3-(TTAGGG)3 (green). DNA was stained with DAPI (magenta). White and green arrows highlight chromatid- and chromosome type fusions respectively. See also figure S1a **b-c.** Quantification of the percentage of telomeres involved in chromatid (b) or chromosome (c) fusions per metaphases after removal of Apollo and/or RAP1 as indicated in the presence or absence of LIG4. Data from 3 independent experiments, 10 metaphases per experiment (n = 30 total), with median. **d.** Representative FISH as in (a) but with metaphase spreads from p53-/-RAP1+/+ or p53-/-RAP1-/-RPE-1 cells as indicated 120 h after transduction with Cas9 and control or Apollo-targeting sgRNA as indicated. **e.** Quantification of the percentage of telomeres fused per metaphases after removal of Apollo as described in (d). Data from 3 independent experiments, 10 metaphases per experiment (n = 30 total), with median. See also, figure S1. All statistics by ordinary One-way ANOVA. ‘ns’ not significant, **P ≤ 0.01 ***P ≤ 0.001 ****P ≤ 0.0001.

We focused out attention on the protective role of RAP1. As shelterin coprecipitates with KU and DNA-PKcs in cell extracts^3,4,24–27^, we considered whether RAP1 might prevent cNHEJ by binding directly to DNA-PK. To test this idea, we used DNase I footprinting to examine the position of purified shelterin and DNA-PK on a blunt-ended telomeric template (Fig. 2a, Fig. S2a). DNA-PK protected 32 bp from the telomere end, consistent with the assembly of a terminally positioned complex^28^ (Fig. 2b, compare lanes 1 and 2). Addition of shelterin reduced the overall efficiency of DNase I cleavage (compare lanes 1 and 6). However, a novel footprint of precisely 10 bp not observed with shelterin alone was also visible directly next to DNA-PK (‘shelterin’ region). A complex of RAP1 and TRF2^29^, but not each component alone was sufficient for this effect (Figs. 2c-e), which was specific to telomeric DNA (Fig. 2f) and observed over a range of protein concentrations (Fig. S2b). Fig. S2c shows the same footprint next to DNA-PK was also observed in the presence of a 3’ overhang (lanes 6 and 8), but only in the absence of POT1/TPP1, which otherwise prevented DNA-PK assembly (lanes 6 and 7).

**Figure 2.**
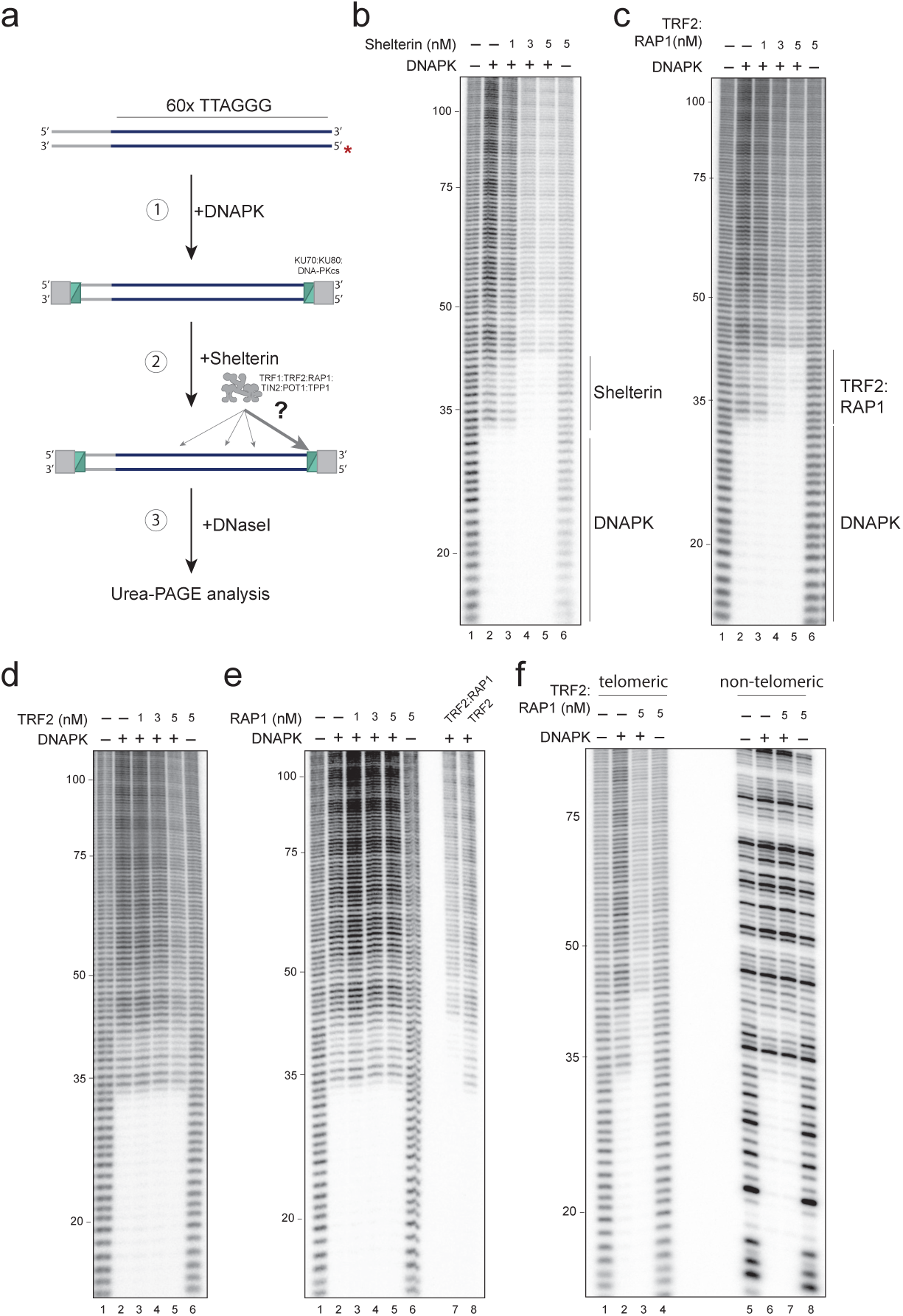
TRF2, RAP1 and DNA-PK form a terminal complex at telomeric DNA ends. **a.** Outline of DNaseI footprinting experiment.^32^P-labelled 5’ end highlighted by a red asterisk. Radiolabelled template is incubated with KU and DNAPKcs prior to the addition of Shelterin composed of TRF1, TRF2, RAP1, TIN2, POT1 and TPP1. DNase I digested products are then analysed by denaturing Urea-PAGE **b-f.** DNaseI footprinting of telomere end-binding complexes. Proteins omitted as indicated. Nucleotides from the 5’ telomeric end indicated.

The data above suggests that DNA-PK bound to telomeric ends can form a sequence-specific complex with TRF2 and RAP1. To examine the architecture of this complex, we determined which domains were required for the DNA-PK-proximal footprint (Fig. 3a, Figs. S2d and e). Binding of RAP1 to TRF2 via the RCT/RBM interface^30^, and binding of TRF2 to DNA via the myb-but not the basic domain was necessary for the extended footprint, suggesting a model in which RAP1 is recruited to DNA by TRF2 for the complex to form (Figs. 3b and c, Fig. S2f). We tested whether this is the primary function of TRF2 in the assay by fusing RAP1 to the DNA-binding domain of fission yeast Teb1, which recognises TTAGGG repeats^15^. In the presence of DNA-PK, the Teb1:RAP1 fusion protein generated a 10 bp footprint indistinguishable from that of RAP1 in complex with TRF2 and prevented by cleavage of the Teb1:RAP1 linker (Fig. 3d and Fig. S2g and h). Thus, the role of TRF2 in forming the complex is to recruit RAP1 to DNA, and the extended footprint is caused by RAP1, not TRF2.

**Figure 3.**
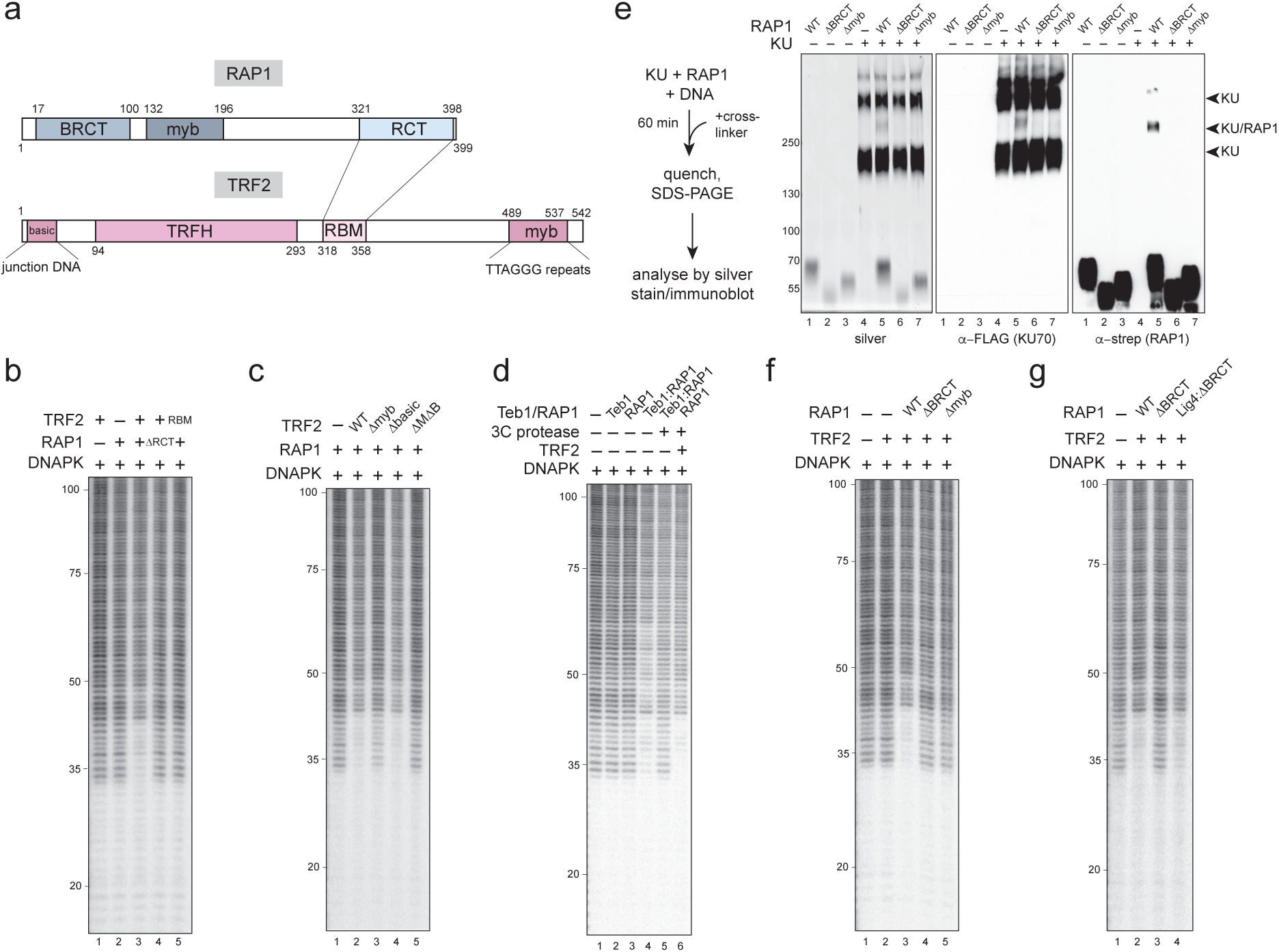
Three distinct interfaces are required for the complex with DNA-PK. **a.** domain organisa-tion of RAP1 and TRF2. **b-d.** DNaseI footprinting of telomere end-binding complexes. Proteins omitted as indicated. See figure S1 for mutant details. **e.** protein cross-linking analysis of RAP1 and KU in the presence of DNA. Proteins were mixed with crosslinker and the products of the reaction were separat-ed on a denaturing tris-acetate polyacrylamide gel and analysed by silver staining or immunoblotting as indicated. Arroheads mark the position of crosslinked species containing only KU, or KU and RAP1 as indicated **f, g.** as in b.

As RAP1 has been proposed to bind KU in cell extracts^25,26^, we examined whether purified RAP1 and KU could bind one another in pulldown assays but did not detect an interaction. However, when RAP1 and KU were mixed in the presence of an amine-specific crosslinker and DNA, we observed a crosslinked species on silver-stained polyacrylamide gels that contained RAP1 and KU as demonstrated by immuno-blotting, suggesting RAP1 mediates a weak interaction with KU (Fig. 3e). Deleting either the RAP1 BRCT- or myb domains disrupted this binding (Fig. 3e) and prevented the extended signal in a footprinting assay (Fig. 3f, compare lanes 3-5). To test whether the BRCT domain could be substituted for a different DNA-PK-binding peptide, we fused RAP1ΔBRCT to the first BRCT domain of LIG4, which recognises KU70^31,32^. Figure 3g shows that adding the LIG4 BRCT domain to RAP1ΔBRCT rescued the 10 bp footprint next to DNA-PK (compare lanes 2-4), confirming that the necessary function of the RAP1 BRCT domain in forming the complex is recognition of KU.

While it is possible that one or other of these regions directly protects the 10 bp next to DNA-PK (see below), these data reveal that three molecular interfaces are required for TRF2 and RAP1 to form a complex with DNA-PK: binding of TRF2 to DNA, binding of TRF2 to RAP1, and binding of RAP1 to KU. The requirement for TRF2 to bind DNA explains why the extended footprint is not formed at non-telomeric ends (Fig. 2f), which do not contain a TRF2 binding site.

To determine how this complex might inhibit cNHEJ at telomeres (figure 1), we used cryo-electron microscopy (cryo-EM) to examine the structure of TRF2, RAP1 and DNA-PK on telomeric DNA. Consecutive rounds of classification followed by focussed refinement yielded an overall structure at 3.58 Å (Fig. 4a, Fig. S3 and Supplementary Video 1). Our map did not contain TRF2, consistent with the notion that its primary role in the complex is to recruit RAP1 to DNA rather than stably bind DNA-PK. The conformation of DNA-PKcs was consistent with previous structures of the unphosphorylated enzyme^33^ (Fig. S4).

**Figure 4.**
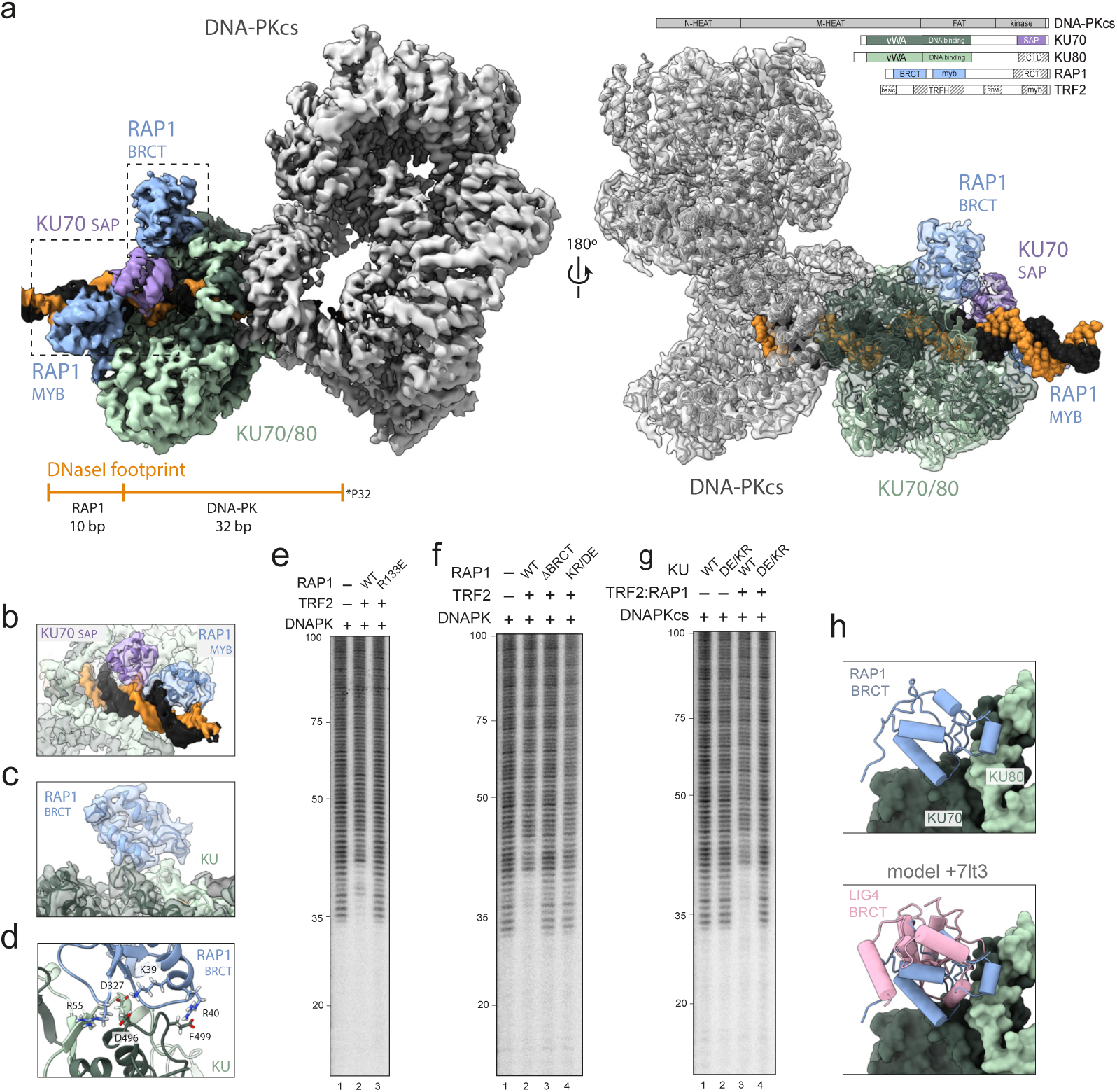
Cryo-EM structure of the RAP1:DNA-PK complex. **a.** Composite electron density map with protein domains coloured as indicated. Schematic shows proteins used for structure determination. Hatching indicates domains not visualised **b.** Subsection of the structure in a, showing the KU70 SAP and RAP1 myb domains bound to DNA. **c. d.** The RAP1 BRCT domain pictured bound to the KU70 vWA domain, highlighting RAP1 and KU residues that mediate the interaction. **e-g.** DNase I footprinting analysis of telomere end-binding complexes with the proteins indicated. RAP1 KR/DE contains mutations K39D, R40E and R55E. KU DE/KR contains mutations KU70 D496K, E499R and KU80 D327K **h.** RAP1 BRCT KU cryo-EM model overlayed with the LIG4 BRCT domain from cryo-EM structure PDB-7It3

Three additional regions of density were observed beyond the core DNA-PK complex: the first sits on the DNA as it enters KU and was modelled as the KU70 SAP domain, visualised here on DNA for the first time (Fig. 4a and b, Fig. S5a). The SAP domain, which remains functionally enigmatic, is positioned on the minor groove with K575, K595 and K596, which have been suggested to bind DNA^34^, coordinating the phosphate backbone (Fig. S5a). The second region engages the neighbouring 10 bp that are protected by RAP1 in our footprinting assays and was modelled as the RAP1 myb domain (Figs. 4a and b, Fig. S5b). The myb adopts a canonical homeodomain arrangement with the recognition helix inserted into the major groove and a conserved N-terminal arm^35^ formed by R133 reaching into the minor groove bound by the KU70 SAP (Fig. 4b, Fig. S5b-d). Chemical crosslinking mass spectrometry confirms the proximity of the myb and SAP regions (Fig. S5e and f). This density is surprising because human RAP1 does not appreciably bind DNA in isolation due to a lack of surface-exposed positive charge^29,36,37^. Our structure suggests DNA-PK complements this deficit by anchoring the myb domain close to DNA through a neighbouring loop bound to the side of KU80 (Fig. S5g). The KU70 SAP is proposed to bind other homeodomain proteins^38^ and additional contacts with RAP1 may also be in place but are not resolved in our structure. A single point mutation in the N-terminal arm (R133E) blocked the footprint next to DNA-PK (Fig. 4e), consistent with the myb:DNA interaction protecting this region.

The third region sits on the KU70 vWA domain and was modelled as the BRCT domain of RAP1 (Figs. 4a and c), positioning a conserved basic patch on RAP1 next to an acidic patch on KU (Fig. 4d and Fig. S6a). Consistent with our structure, charge reversal mutations in these regions prevented the extended signal in a footprinting assay (Fig. 4f and g, RAP1 KR/DE and KU DE/KR) and crosslinking of RAP1 to KU (Fig. S6c compare lanes 7, 9 and 11). Remarkably, overlaying the RAP1 BRCT domain with the LIG4 BRCT domain bound to KU^32^ shows that RAP1 at this position directly occludes the binding site for LIG4 (Fig. 4h). These data suggest that by forming a complex with DNA-PK, TRF2 and RAP1 may prevent cNHEJ by simply blocking Ligase 4 recruitment.

To test this idea, we developed a pulldown assay in which purified KU was immunoprecipitated after incubation with purified DNA-PKcs, purified XRCC4/LIG4 and a short DNA template (Fig. 5a). DNA-PKcs associated with KU in a DNA-dependent manner, consistent with the assembly of DNA-PK at a DNA end (Fig. S7a). XRCC4/LIG4 recruitment was sensitive to mutations in the acidic patch on KU that is required to bind RAP1 and known to bind LIG4^32^ (Fig. S7a, compare lanes 7 and 8). While TRF2 or RAP1 had a negligible effect on the assay individually, recruitment of XRCC4/LIG4 was blocked when they were added together, even when XRCC4/LIG4 was preincubated with DNA-PK and present in large excess (Fig. 5b and Figs. S7b and c). To test whether the ability of TRF2/RAP1 to block LIG4 recruitment required binding to DNA-PK, we repeated the experiment with RAP1 ΔBRCT or KR/DE, which are unable to bind KU (Figure 4). Inhibition of LIG4 recruitment was not observed under these conditions but could be restored by fusing the N-terminal BRCT domain of LIG4 onto the N-terminus of RAP1ΔBRCT (Fig. 5c and Fig. S7d). Figure 5d shows RAP1 R133E was also defective in preventing the recruitment of LIG4, suggesting multiple contacts between RAP1, KU and DNA are required for this effect.

**Figure 5.**
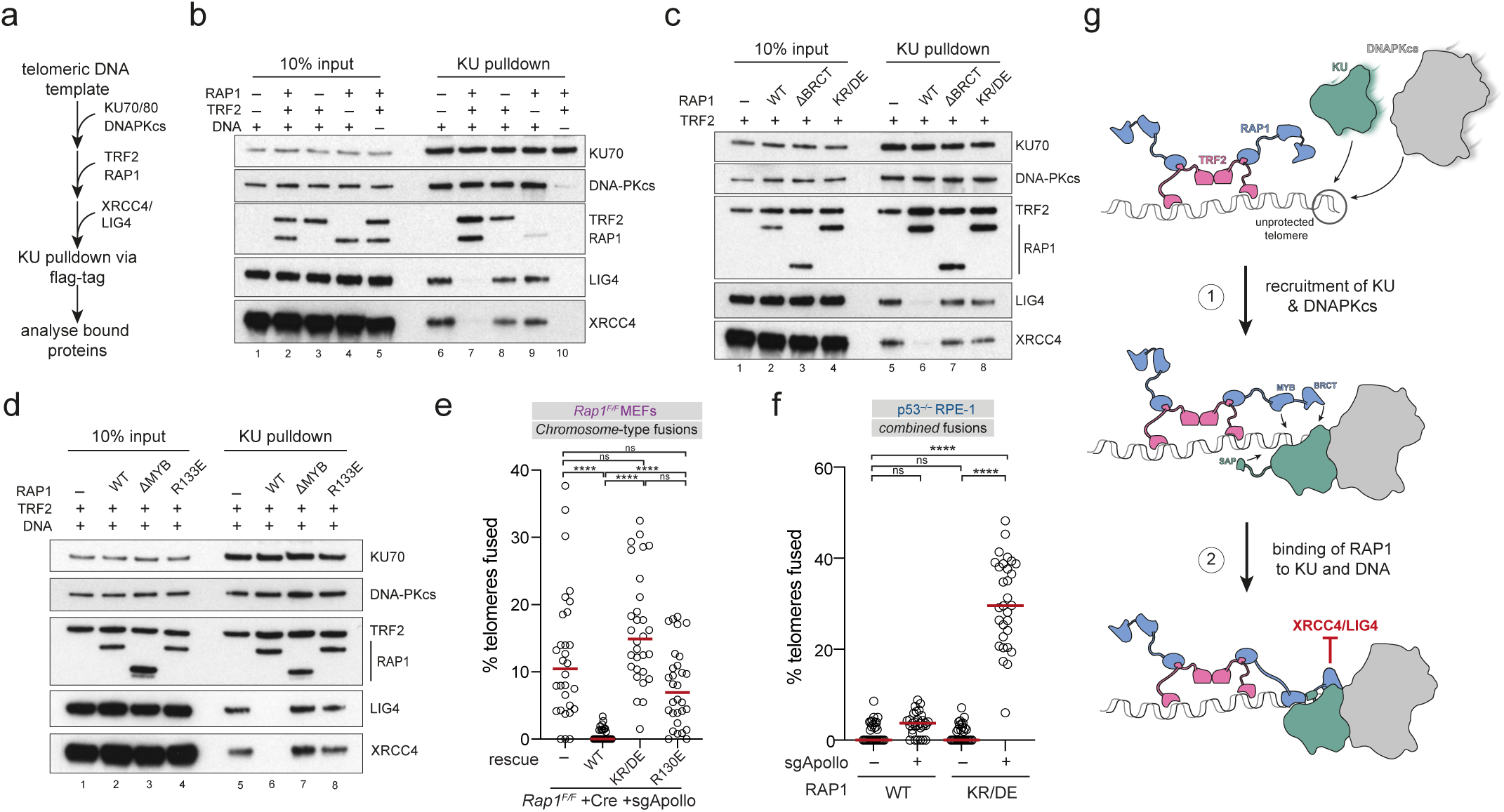
TRF2 and RAP1 directly prevent recruitment of XRCC4/LIG4 to DNA-PK. **a.** Outline of the KU pulldown approach. See methods for details **b-d.** KU-bound proteins from reactions containing KU70/80, DNA-PKcs, XRCC4/LIG4, TRF2, RAP1 and template DNA unless indicated were separated by SDS-PAGE and immunoblotted as indicated. TRF2, RAP1 and LIG4 detected with anti-strep tag antibody, KU70 with anti-FLAG antibody. As shown in (b), associated of TRF2 with KU is mediated by template DNA **e.** Quantification of the percentage of telomeres involved in chromosome fusions per metaphase upon over-expression of mouse RAP1, RAP1 KR/DE and RAP1 R130E after Crispr- and Cre-mediated deletion of Rap1 and Apollo respectively in ApolloF/F Lig4+/+ MEFs. Data from 3 independent experiments, 10 metaphases per experiment (n = 30 total), with median. Statistics by ordinary One-way ANOVA. ‘ns’ not significant, ****P ≤ 0.0001. See Fig. S8 for further details. **f.** Quantification of the percentage of telomeres fused per metaphase upon Crispr-mediated deletion of Apollo in p53-/-RPE-1 cells with wildtype RAP1 or the RAP1 KR/DE mutant. Data from 3 independent experiments, 10 metaphases per experiment (n = 30 total), with median. Statistics as in (e). See Fig. S8 for further details **g.** Model for the direct inhibition of DNA-PK by TRF2/RAP1 at mammalian telomeres. When assembled on telomeric DNA, DNA-PK is bound by the RAP1 myb and BRCT domains, which acts as a circuit breaker that prevents DNA-PK from engaging LIG4.

To examine whether inhibition of LIG4 recruitment is required for RAP1 to prevent cNHEJ at telomeres, *rap1-/-apollo-/-* MEFs were complemented with the RAP1 mutants examined above (Fig 5e and Fig. S8a-c). Figure 5e shows RAP1 that is unable to form a complex with DNA-PK and block LIG4 recruitment could not protect telomeres from cNHEJ (Fig. S8c). Figure 5f demonstrates this is also the case in human RPE-1 cells (Fig. 5f and Fig. S8d-g).

RAP1 is the most highly conserved component of eukaryotic chromosome ends, yet its role at mammalian telomeres has remained elusive. Our results reveal it is the defining component of an inhibitory pathway coordinated by TRF2 in which RAP1 directly and specifically supresses the end joining function of DNA-PK by preventing the recruitment of Ligase 4 (Fig. 5g). These findings provide a molecular basis for previous studies implicating human RAP1 in chromosome end protection^15–18^, offer a mechanism for how cNHEJ can be blocked at telomeres with diverse end structures or lacking t-loops^13,39^, and resolve the long-standing paradox that DNA-PK can bind to telomeres and regulate their resection without activating cNHEJ^3,4^. By demonstrating that the assembly of DNA-PK can be functionally uncoupled from its end joining activity through LIG4 recruitment, our study also expands the mechanisms available to regulate pathway choice during double strand break repair.

Apollo and RAP1 are recruited to telomeres via TRF2, which therefore acts as a master regulator of two equally effective pathways to block cNHEJ. Why is RAP1 employed when Apollo is apparently sufficient? Processing of telomeres by Apollo depends on DNA-PK^23^ and a TRF2 binding motif specific to vertebrate homologues^40^. As the RAP1:KU interaction is broadly conserved across metazoa (Fig. S9), we propose the pathway described here predates Apollo’s telomeric function and enabled DNA-PK at leading strand telomeres to be coopted into a resection role by preventing it from activating cNHEJ. In vertebrate cells, RAP1 will ensure DNA-PK on blunt leading strand ends cannot engage cNHEJ either before resection occurs or in instances where resection fails. There may also be scenarios in which RAP1 is the primary protective factor. For example, RAP1 deletion alone increases cNHEJ at telomeres in senescent cells^16,41^. Given that only a small number of telomeric repeats are required for TRF2 and RAP1 to prevent LIG4 recruitment in vitro (Fig. 5b), the mechanism described here may be adept at protecting critically short telomeres in these cells from cNHEJ.

## Methods

### DNA templates

A 2.8 kb plasmid containing 360 bp of telomeric DNA was amplified in SURE2 *E. coli* cells grown at 30°C and extracted using a QIAGEN Plasmid Maxi kit. The plasmid was linearised by BsmFI digestion for 1 hour at 37°C leaving one end with telomeric TTAGGG repeats. The DNA was dephosphorylated with Quick-CIP for 30 min at 37°C and cleaned up by phenol chloroform extraction and ethanol precipitation. The DNA was subsequently phosphorylated using PNK and [γ-32P]ATP for 1 hour at 37°C, passed over a G-50 desalting column and phenol chloroform extracted. Labelled DNA was digested by SacI for 1 hour at 37°C to isolate a 390-bp DNA fragment ending in 60 TTAGGG-repeats from the rest of the plasmid. Digested DNA was run on a 10% TBE-PAGE gel (Invitrogen) after which the telomeric fragment was excised and gel extracted by shaking in 10 mM Tris pH 8.0, 300 mM NaCl, 1 mM EDTA overnight at 21°C. The final DNA fragment (PE1, see table 1 for sequence) was isopropanol precipitated and resuspended in 1x TE buffer. A non-telomeric DNA fragment (PE2, see table 1 for sequence) containing 360 bp of random DNA sequence in place of TTAGGG repeats was also prepared through the same procedure using a non-telomeric plasmid. Telomeric DNA substrates for crosslinking experiments were prepared by mixing oligonucleotides PE3 and PE4 at a 1:1 molar ratio, heating to 95°C and slowly cooling to room temperature over 2 hours.

**Table 1.**
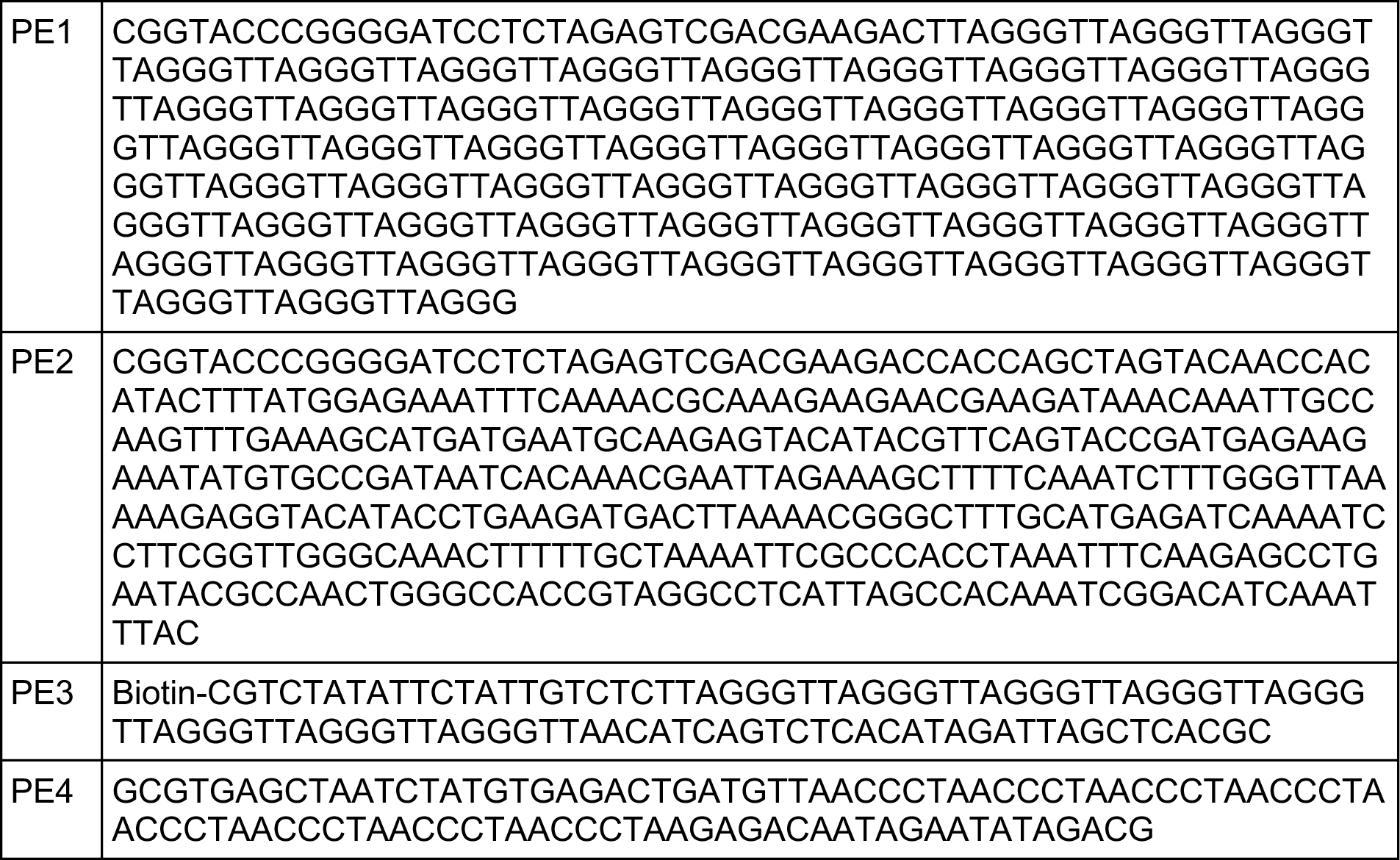
Oligonucleotides used.

### Protein expression

Open reading frames for human KU70/80, RAP1 or TRF2 were codon optimised for *S. frigiperda* and cloned into pACEBAC1 vector. The mutants indicated were generated using PCR-based mutagenesis (see table 2 for mutant details). The shelterin genes (TERF1, TERF2, TERF2IP/RAP1, TINF2, ACD/TPP1, POT1), codon-optimised for *E. coli*, were synthesised by GenScript and cloned into pACEBAC1, using a nicking cloning system^42^. Vectors were transposed into DH10 MultiBac Competent *E. Coli* cells and grown in LB medium overnight at 37°C shaking at 200 rpm. Bacmid DNA was extracted and used to transfect Sf9 insect cells which were subsequently grown at 27°C, shaking at 130 rpm over several cell passages. At passage three (P3), 200 ml of High Five insect cells at a 1.5 x 10^6^ density were infected with baculoviruses for protein expression, incubating at 27°C with 130 rpm. After 3 days, the cell count was checked, and cells were harvested by centrifugation at 760 x g for 20 minutes at 4°C. Cell pellets were then resuspended in PBS, transferred into 50-ml Falcon tubes and pelleted again by centrifugation at 470 x g for 25 minutes. Supernatants were discarded, the pellets flash frozen in liquid nitrogen and placed at −80°C until required.

**Table 2.**
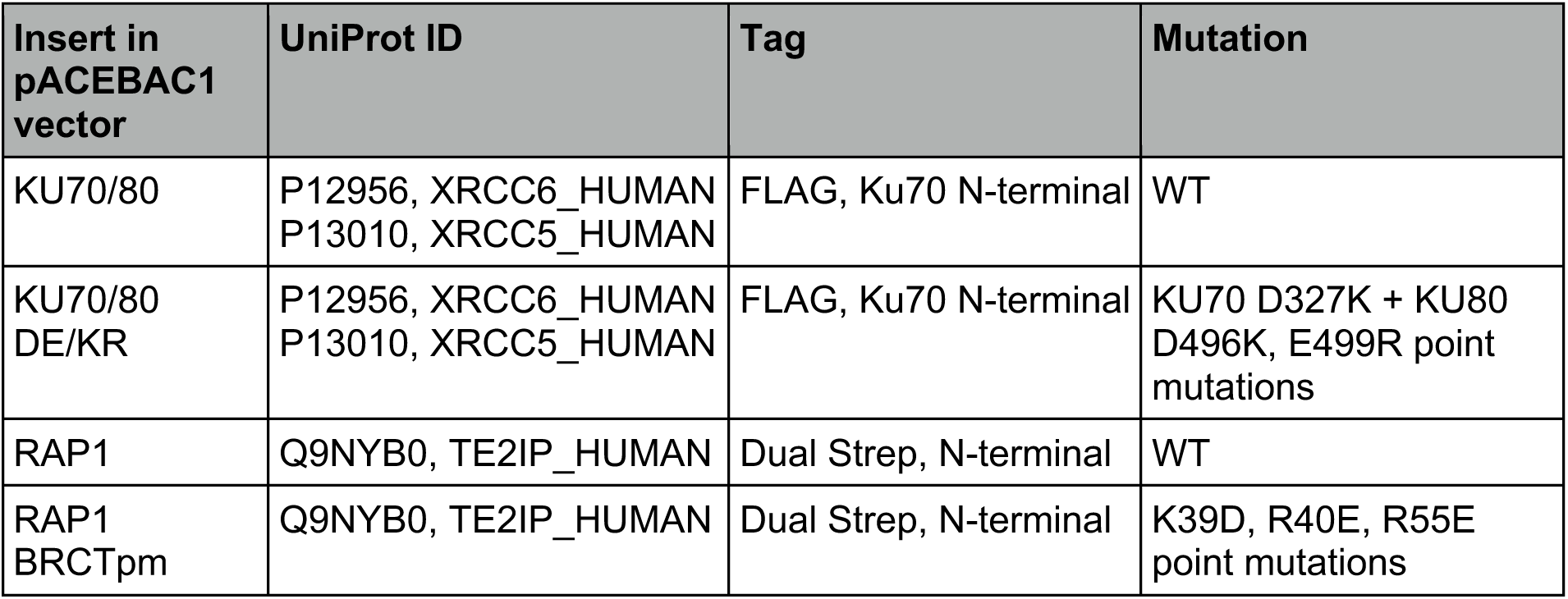
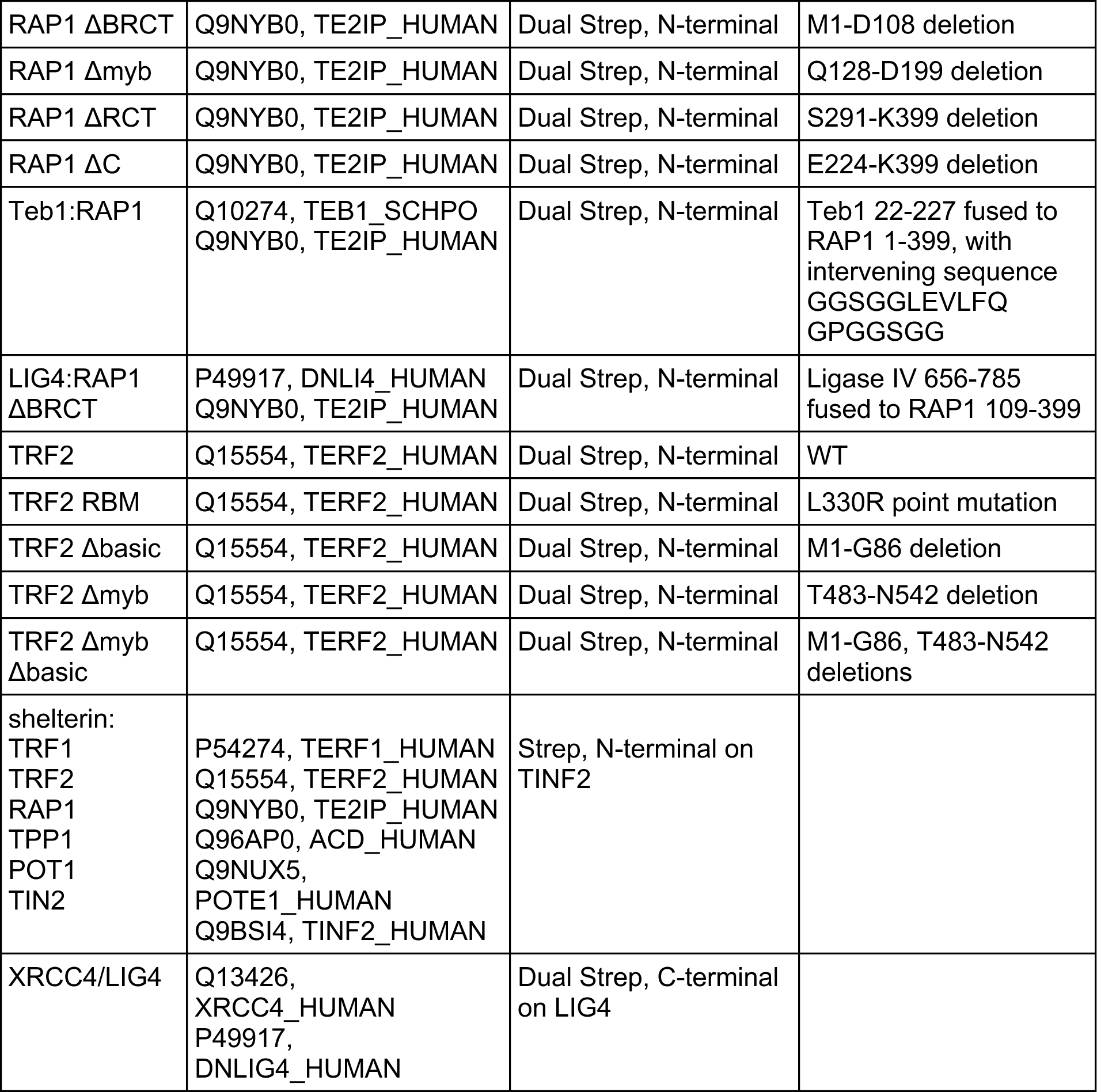
Constructs used.

### Nuclear extract preparation

HeLa cell pellets were resuspended in buffer A (10 mM HEPES pH 8.0, 10 mM KCl, 1.5 mM MgCl_2_, 0.5 mM DTT, 0.5 mM AEBSF) and incubated for 10 min at 4°C. Following centrifugation at 1,033 x g for 10 min at 4°C the pellet was resuspended in two pellet volumes of buffer A and lysed by 20 strokes in a Dounce homogeniser (pestle type B). Lysed cells were centrifuged at 25,000 x g for 20 min at 4°C and the pellet was resuspended in 1.3x pellet volumes of buffer C (20 mM HEPES pH 8.0, 420 mM NaCl, 1.5 mM MgCl_2_, 0.2 mM EDTA, 25 % glycerol, 0.5 mM DTT, 0.5 mM AEBSF). Cells were resuspended by 20 strokes in a Dounce homogeniser (pestle type B), incubated for 30 min at 4°C and centrifuged at 25,000 x g for 30 min at 4°C. The supernatant was collected, and flash frozen in liquid nitrogen.

### Protein purification

**DNA-PKcs purification.** HeLa cell nuclear extract was diluted into DPKQ buffer (20 mM HEPES ph 7.6, 100 mM NaCl, 2 mM MgCl2, 0.5 mM EDTA, 10% glycerol, 0.5 mM DTT, 0.5 mM AEBSF) and ultracentrifuged at 50,000 x g for 1 hour at 4°C. The supernatant was collected and filtered through a 0.45 µm syringe filter before injection onto a Q-sepharose column equilibrated in DPKQ buffer. The column was washed in DPKQ buffer and proteins were eluted using a 100 mM - 1 M NaCl gradient. Fractions were spotted onto a nitrocellulose membrane and western blotted, probing for DNA-PKcs. Fractions with DNA-PKcs were pooled, diluted into DPKQ buffer and loaded onto a heparin column equilibrated in the same buffer. The column was washed DPKQ buffer and proteins were eluted using a 100 mM - 1 M NaCl gradient. DNA-PKcs fractions were pooled, dialysed into DPKC buffer (50 mM Tris pH 7.3, 0.5 mM EDTA, 5% glycerol, 2.5 mM DTT, 0.5 mM AEBSF) with 50 mM KCl, and loaded onto a dsDNA-conjugated CNBr-activated Sepharose column equilibrated in the same buffer. The column was washed in DPKC buffer with 50 mM KCl and bound proteins were eluted in DPKC buffer with 411 mM KCl. DNA-PKcs fractions were pooled, diluted into DPKC buffer with 100 mM KCl and 0.02% Tween-20 and loaded onto a MonoQ column equilibrated in the same buffer. The column was washed in DPKC buffer with 100 mM KCl and 0.02% Tween-20 and proteins were eluted using a 100 mM - 1 M KCl gradient. DNA-PKcs fractions were pooled, diluted into DPKC buffer with 100 mM KCl and 0.02% Tween-20 and loaded onto a MonoS column equilibrated in the same buffer. The column was washed in DPKC buffer with 100 mM KCl and 0.02% Tween-20 and proteins were eluted using a 100 mM - 1 M KCl gradient. Final DNA-PKcs fractions were pooled, concentrated using a 100 kDa Amicon Ultra centrifugal filter, flash frozen in liquid nitrogen and stored at −80 °C. **KU70/80 purification.** Cell pellets were resuspended in 50 mM Tris pH 8.0, 2 mM β-ME with protease inhibitor tablets and incubated stirring for 20 min at 4 °C. Cells were lysed by the addition of 16.7% glycerol and 300 mM NaCl, stirring for 30 min at 4°C. Lysed cells were ultracentrifuged at 125,000 x g for 30 min at 4°C and the supernatant was incubated with anti-FLAG resin for 2 hours at 4 °C. Beads were successively washed in Ku buffer (50 mM Tris pH 8.0, 5% glycerol, 2 mM β-ME) with 300 mM NaCl followed by Ku buffer with 150 mM NaCl. Proteins were eluted using the latter buffer supplemented with 3x FLAG peptide. Proteins were subsequently separated on a Superdex 200 gel filtration column, equilibrated and run using Ku buffer with 150 mM NaCl. Final Ku70/80 fractions were pooled, concentrated using a 30 kDa Amicon Ultra centrifugal filter, flash frozen in liquid nitrogen and stored at −80 °C. **TRF2, RAP1 and shelterin purification.** For TRF2 and RAP1, cell pellets were resuspended in 50 mM HEPES pH 7.6, 1 mM DTT with EDTA-free protease inhibitor tablets (one per 50 ml - Roche) and incubated with stirring for 20 min at 4 °C. Protein was extracted by the addition of 16.7% glycerol and 300 mM NaCl, stirring for 30 min at 4°C. Extract was cleared by ultracentrifugation at 125,000 x g for 30 min at 4 °C. The supernatant was applied to Strep-Tactin XT 4Flow resin equilibrated in shelterin buffer (50 mM HEPES pH 7.6, 500 mM NaCl, 10% glycerol, 1 mM DTT). Beads were washed in shelterin buffer and proteins were eluted with the same buffer supplemented with 10 mM biotin. Proteins were subsequently separated on a Superdex 200 10/300 column equilibrated and run in shelterin buffer. Final TRF2 or RAP1 fractions were pooled, concentrated using a 30 kDa Amicon Ultra centrifugal filter, flash frozen in liquid nitrogen and stored at −80 °C. To cleave the TEB1:RAP1 fusion protein, 0.24 µM PreScission protease was incubated with 1.2 µM TEB1:RAP1 for 2 hours at 4 °C prior to experiments. For the shelterin complex, cell pellets were resuspended in lysis buffer (50 mM Hepes pH 8.0, 300 mM NaCl, 10 % glycerol, 1 mM MgCl_2_, 10 mM beta-mercaptoethanol, 0.1 µl/ml Base muncher nuclease, 8 µg/ml Avidin, 1 mM AEBSF and EDTA-free protease inhibitor tablets (one per 50 ml - Roche)) and cell were lysed by sonication. Lysate was cleared by centrifugation at 48,380 x g for 1 h, 4 °C. Cleared lysate was passed through a 0.45 µm filter and applied to a StrepTrap column equilibrated with StrepTrap wash buffer (50 mM Hepes 8.0, 300 mM NaCl, 10 glycerol and 1 mM TCEP), which was washed with 20 CV StrepTrap wash buffer prior to elution with 5 CV StrepTrap wash buffer supplemented with 10 mM d-Desthiobiotin. 0.5 ml of eluate containing the highest concentration of shelterin was applied to a Superose 6 10/300 column preequilibrated in 50 mM Hepes pH 8.0, 300 mM NaCl, 10 % glycerol and 1 mM TCEP, and run in the same buffer. The desired fractions were aliquoted and flash frozen for storage. Further characterisation of the full shelterin complex will be reported elsewhere. **XRCC4/LIG4 purification.** Cell pellets were resuspended in 50 mM HEPES pH 7.6, 1 mM DTT, 1mM EDTA with EDTA-free protease inhibitor tablets (Roche – 1 per 50 ml buffer) and incubated with stirring for 20 min at 4°C. Proteins were extracted by the addition of glycerol to 16.7% and NaCl to 300 mM with stirring for 30 min at 4°C. Extract was centrifuged at 41,656 x g for 30 min at 4°C and the supernatant was applied to Strep-Tactin XT 4Flow resin equilibrated in X4 buffer (50 mM HEPES pH 7.6, 300 mM NaCl, 10 % glycerol, 1 mM DTT, 1mM EDTA). Beads were washed in X4 buffer and proteins were eluted with the same buffer supplemented with 30 mM biotin. Protein fractions were pooled, the NaCl concentration diluted to 100mM then applied to a 1 ml heparin column and subjected to a linear gradient from 0.1 M to 1 M NaCl over 20 column volumes. Proteins were subsequently separated on a Superdex 200 gel filtration column equilibrated and run in 100mM NaCl buffer (50 mM HEPES pH 7.6, 100 mM NaCl, 10% glycerol, 1 mM DTT, 1mM EDTA). Final XRCC4/LIG4 fractions were pooled, concentrated using a 30 kDa Amicon Ultra centrifugal filter, flash frozen in liquid nitrogen and stored at −80 °C.

### Nano differential scanning fluorimetry

Purified proteins as indicated were analysed by nano differential scanning fluorometry to assess thermal protein stability using a Tycho NT.6 (Nanotemper) with 10 µl capillaries, monitoring fluorescence at 330 and 350 nm over a 35-95°C temperature ramp (30 °C/min).

### DNase I footprinting

2 nM [γ-^32^P]-PNK-labelled PE1 (telomeric) or PE2 (non-telomeric) template was mixed with 30 nM DNA-PKcs and 50 nM KU70/80 in 25 mM HEPES pH 7.6, 80 mM KCl, 1.5 mM CaCl_2_, 1.5 mM MgCl_2_, 5% glycerol, 50 µg/ml BSA, 2 mM DTT and incubated on ice for 5 min. 5 nM shelterin or TRF2:RAP1 (dimer concentration of TRF2) was added to DNA-bound DNA-PK and incubated at 37 °C for 10 min. Nuclease cleavage was initiated by addition of DNaseI to 0.5 U/ml and the reactions were incubated for a further 2 min at 37°C before quenching with 25 mM EDTA, 0.2% SDS, 0.2 mg/ml Proteinase K. Following incubation at 37 °C for 10 min, samples were extracted with phenol chloroform, ethanol precipitated and resuspended in 2 µl 99% formamide, 5 mM EDTA, bromophenol blue. Samples were boiled for 2 min and run on an 8% urea-PAGE sequencing gel in 1x TBE.

### Electrophoretic Mobility Shift Assay

1 nM [γ-^32^P]-labelled telomeric DNA was mixed with 2, 10 or 40 nM TRF2 or RAP1 in 25 mM HEPES ph 7.6, 80 mM KCl, 1.5 mM CaCl_2_, 1.5 mM MgCl_2_, 5% glycerol, 50 µg/ml BSA, 2 mM DTT. 10 µl reactions were incubated at 37°C for 15 min. Samples were supplemented with 1% sucrose, Orange G and run on a 1.5 % agarose gel in 0.5x TBE.

### Crosslinking

200 nM KU70/80 was mixed with 200 nM RAP1 in 20 mM HEPES pH 7.6, 200 nM NaCl, 2 mM MgCl_2_, 0.5 mM EDTA, 10% glycerol, 0.5 mM DTT, 0.5 mM AEBSF and incubated for 5 min at 4°C. Samples were supplemented by 100 nM annealed DNA substrates and incubated for 10 min at 4 °C. Proteins were crosslinked by addition of 2 mM DSSO and incubated for 60 min at room temperature. Reactions were stopped with 20 mM Tris pH 7.6. Samples were run on a Criterion XT 3-8% Tris-Acetate PAGE gel in 1x XT Tricine and analysed by silver staining (SilverQuest, Invitrogen) or immunoblotting, probing for KU70 or RAP1.

### DNA-PK Pulldown

10 nM KU70/80 and 15 nM DNA-PKcs were incubated with 5 nM annealed DNA oligos (see *‘DNA templates’*) for 3 min at 30°C in 25 mM HEPES pH 7.6, 80 mM KCl, 10 % glycerol, 0.02 % NP-40 and 1 mM DTT in a final volume of 25 µl. 5 nM TRF2:RAP1 complex was added, and after a further 3 min, 30 nM XRCC4:LIG4 complex was added. After a further 3 min, the complete reaction was added to 1 µl equivalent of anti-FLAG M2 magnetic beads (Sigma), and the mixture incubated at 4 °C with shaking for 30 min. Beads were pelleted on a magnetic rack, washed 3x with 50 µl 25 mM HEPES pH 7.6, 80 mM KCl, 10 % glycerol, 0.02 % NP-40 and 1 mM DTT with a brief vortex included for each wash. Beads were resuspended in wash buffer supplemented with 0.25 mg/ml 3x FLAG peptide and incubated for 20 min at 18 °C with shaking. Eluted proteins were supplemented with SDS loading buffer, run on a 4-12 % TGX precast gel (Biorad), proteins transferred onto nitrocellulose membrane at 80 V for 90 min and detected by immunoblotting with the antibodies indicated.

### Cryo-EM sample preparation

Annealed PE03 and PE04 (see ‘DNA templates’) were mixed 1:1 with streptavidin in TE buffer and incubated for 30 min at room temperature. Streptavidin-bound DNA was diluted to 250 nM in 40 mM HEPES pH 7.6, 100 mM NaCl, 3 mM MgCl2, 1 mM DTT, 2.5% glycerol and incubated with 250 nM KU70/80 and 250 nM DNA-PKcs for 10 min at 4 °C. 250 nM TRF2:RAP1 (dimer concentration of TRF2) was added and incubated for another 5 min at 4°C. Samples were supplemented with 0.05% CHAPS prior to cryo-EM grid preparation.

### Cryo-EM data acquisition and image processing

300-mesh copper R1.2/1.3 grids (Quantifoil) were coated with a thin layer or carbon and glow-discharged at 15 mA for 30 s (PELCO easiGlow). 3 µl sample was applied to glow-discharged grids and incubated for 5 s followed by blotting for 3 sec using a Vitrobot Mark IV (FEI ThermoFisher) operated at 4 °C and 100% humidity. Grids were subsequently plunge-frozen in liquid ethane. Cryo-EM data were acquired at 200 kV on a Glacios Cryo-TEM equipped with a Falcon 4i Direct Electron Detector. 30,604 movies with 30 frames were collected at x150,000 magnification (0.94 Å pixel size) with a total electron dose of 50 e^-^/Å^2^ and a defocus range of −1.0 to −2.5 μm. Subsequent image processing was performed in cryoSPARC (v4.3.1)^43^. Movies were motion corrected using patch alignment with all frames followed by patch CTF estimation. Particles were picked through automated template-based picking (template EMD-6803) and extracted with x4 binning using a box size of 96 pixels. Following 2D classification 1,464,362 DNA-PK particles were selected to reconstruct an ab-initio 3D model, subsequently used as a starting model for heterogeneous refinement using 5 classes. The most prominent DNA-PK class was selected and subjected to homogeneous refinement followed by local refinement using a focus mask encompassing the KU70/80:DNA core. 3D classification without alignment using the same mask and 10 classes was performed to identify particles containing additional Rap1 and KU densities. Classes lacking either the RAP1 BRCT, the RAP1 myb or the KU70 SAP domain were excluded. 526,885 selected particles were re-extracted unbinned with a 384-pixel box size and a 3D map was reconstructed through homogeneous refinement. Following global CTF refinement a structure of the full DNA end-binding complex was resolved to 3.58 Å using homogeneous refinement. The KU:RAP1:DNA core was locally refined to 3.32 Å using a focus mask excluding DNA-PKcs. The DNA-PKcs:DNA density was similarly refined using a mask excluding the KU-RAP1-DNA core and subjected to 3D classification without alignment using the same mask. Some flexibility in DNA-PKcs conformation was observed and classes of the most prominent conformation containing 370,172 paricles were selected. A DNA-PKcs:DNA structure from these particles was resolved to 3.40 Å by homogeneous refinement followed by local refinement using a DNA-PKcs focus mask. Maps for the full end-binding complex and the locally refined densities were sharpened in cryoSPARC and combined using Phenix (v1.20.1) combine_focused_maps^44^. The composite map (EMD-19065) was subsequently used for model building in Coot^45^ and figure generation in ChimeraX^46^.

### Model Building and Validation

Molecular models for human DNA-PK (PBD 7K1K), RAP1 myb (PDB 1FEX) and KU70 SAP (PDB 1JJR) were docked into the cryo-EM map using the Fit in Map command in ChimeraX^46^. The RAP1 BRCT domain from a KU:RAP1 Alphafold model (see below) was similarly docked into the cryo-EM map. Models were refined against the map using Namdinator^47^ followed by manual inspection in Coot^45^. Unoccupied protein densities and nucleic acids were modelled de novo. The resulting model was iteratively refined using Phenix (v1.20.1) real_space_refinement^48^ with geometry and secondary structure restraints followed by manual adjustment in Coot. The quality of the final atomic model (PDB 8RD4) was evaluated by MolProbity^49^ (see table 3).

**Table 3.**
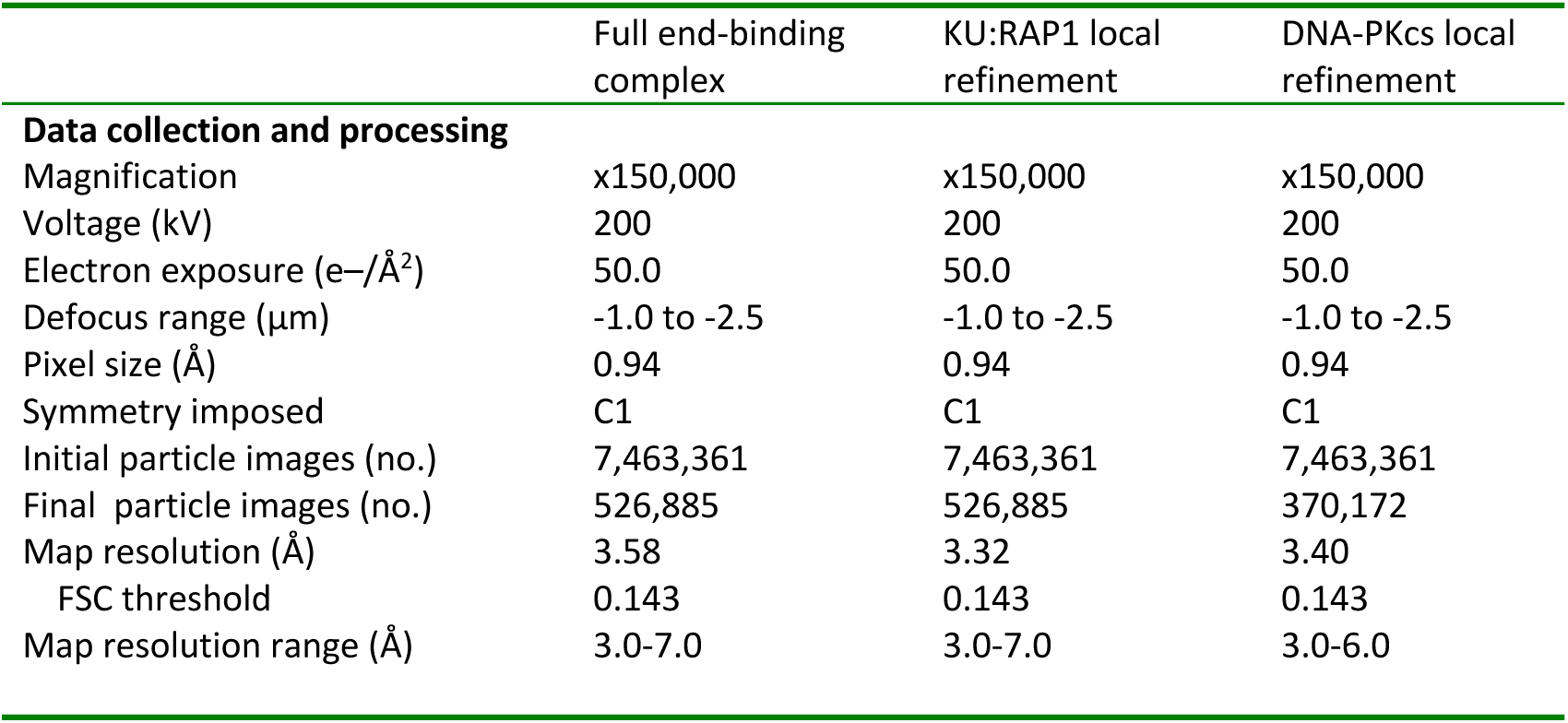
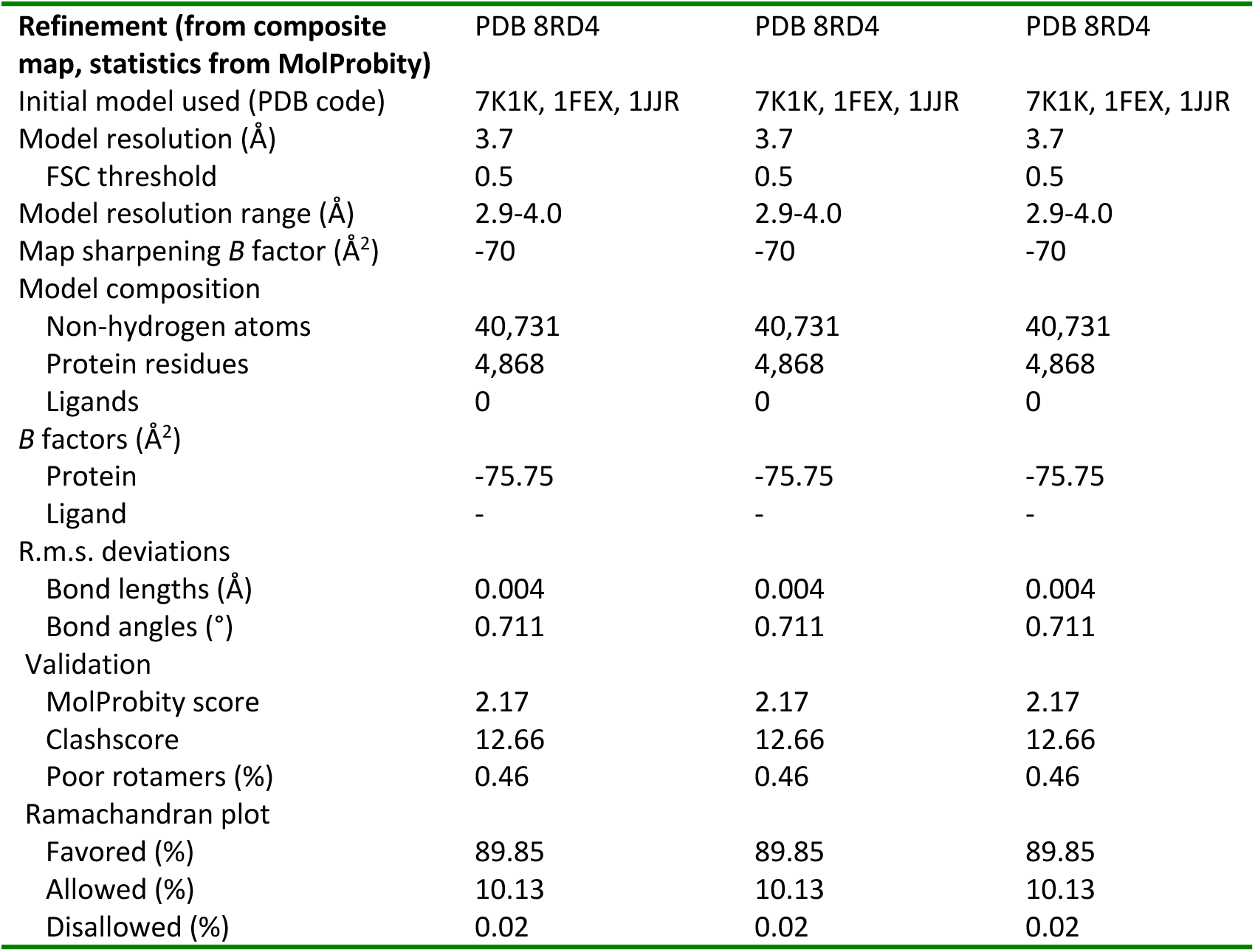
Cryo-EM parameters.

|  | Full end-binding complex | KU:RAP1 local refinement | DNA-PKcs local refinement |
| --- | --- | --- | --- |
| **Data collection and processing** |  |  |  |
| Magnification | x150,000 | x150,000 | x150,000 |
| Voltage (kV) | 200 | 200 | 200 |
| Electron exposure (e-/Å<sup>2</sup>) | 50.0 | 50.0 | 50.0 |
| Defocus range (μm) | -1.0 to -2.5 | -1.0 to -2.5 | -1.0 to -2.5 |
| Pixel size (Å) | 0.94 | 0.94 | 0.94 |
| Symmetry imposed | C1 | C1 | C1 |
| Initial particle images (no.) | 7,463,361 | 7,463,361 | 7,463,361 |
| Final particle images (no.) | 526,885 | 526,885 | 370,172 |
| Map resolution (Å) | 3.58 | 3.32 | 3.40 |
| FSC threshold | 0.143 | 0.143 | 0.143 |
| Map resolution range (Å) | 3.0-7.0 | 3.0-7.0 | 3.0-6.0 |

| **Refinement (from composite map, statistics from MolProbity)** | **PDB 8RD4** | **PDB 8RD4** | **PDB 8RD4** |
| --- | --- | --- | --- |
| Initial model used (PDB code) | 7K1K, 1FEX, 1JJR | 7K1K, 1FEX, 1JJR | 7K1K, 1FEX, 1JJR |
| Model resolution (Å) | 3.7 | 3.7 | 3.7 |
| FSC threshold | 0.5 | 0.5 | 0.5 |
| Model resolution range (Å) | 2.9-4.0 | 2.9-4.0 | 2.9-4.0 |
| Map sharpening *B* factor (Å<sup>2</sup>) | -70 | -70 | -70 |
| Model composition |  |  |  |
| Non-hydrogen atoms | 40,731 | 40,731 | 40,731 |
| Protein residues | 4,868 | 4,868 | 4,868 |
| Ligands | 0 | 0 | 0 |
| *B* factors (Å<sup>2</sup>) |  |  |  |
| Protein | -75.75 | -75.75 | -75.75 |
| Ligand | - | - | - |
| R.m.s. deviations |  |  |  |
| Bond lengths (Å) | 0.004 | 0.004 | 0.004 |
| Bond angles (°) | 0.711 | 0.711 | 0.711 |
| Validation |  |  |  |
| MolProbity score | 2.17 | 2.17 | 2.17 |
| Clashscore | 12.66 | 12.66 | 12.66 |
| Poor rotamers (%) | 0.46 | 0.46 | 0.46 |
| Ramachandran plot |  |  |  |
| Favored (%) | 89.85 | 89.85 | 89.85 |
| Allowed (%) | 10.13 | 10.13 | 10.13 |
| Disallowed (%) | 0.02 | 0.02 | 0.02 |

### Alphafold modelling

Full length human RAP1, KU70 and KU80 were analysed using Alphafold 3 on the online Alphafold server.

### Protein alignments

Pre-computed protein alignments were analysed using the Proviz online tool^50^

### Antibodies used

Human DNA-PKcs was detected with antibody 18-2 (Invitrogen MA5-13238). Recombinant human KU70 was detected via an N-terminal FLAG-tag with antibody M2 (Sigma F1804). Recombinant human LIG4, TRF2 and RAP1 were detected via a dual strep tag using antibody (Abcam ab76949). Recombinant human XRCC4 was detected with antibody C-4 (Santa Cruz sc-271087). Mouse RAP1 was detected with antibody D9H4 (Rabbit mAb #5433), human RAP1 with A300-306A antibody (Bethyl Laboratories) mouse and human TRF2 with D1Y5D (Cell Signaling 13136), and Donkey anti-Rabbit IgG horseradish peroxidase (NA934V, Cytiva) or Goat anti-Rabbit IgG horseradish peroxidase (31460, Invitrogen) secondary antibodies. Mouse β-actin was detected with primary antibody C4 (Cell Signaling #3700) and Goat anti-Mouse IgG horseradish peroxidase (31430, Invitrogen) secondary antibody. Human Apollo was detected with HPA064934 (Atlas Antibodies).

### Cell lines and Viral gene delivery

SV40-LT *Apollo*^F/F^ Lig4^+/+^, *Apollo*^F/F^ Lig4^-/-^, and *Trf2*^F/F^*Rosa26Cre-ER*^T1^ MEFs have been previously described (Wu et al, 2010; Lottersberger et al, 2015). hTERT immortalized RPE-1 cells have been previously described^51^ All MEFs were immortalized with pBabeSV40LargeT and cultured in Dulbecco’s Modified Eagle Medium (DMEM, Cytiva) supplemented with 15% fetal bovine serum (FBS, Gibco), non-essential amino acids (Cytiva), L-glutamine (Cytiva), penicillin/streptomycin (Cytiva), 50 µM β-mercaptoethanol (Sigma). 293T and Phoenix eco cells (ATCC, Rockville, MD) were cultured in DMEM (Cytiva) supplemented with 10% HyClone Bovine Calf Serum (Cytiva), non-essential amino acids (Cytiva), L-glutamine (Cytiva), and penicillin/streptomycin (Cytiva). RPE-1 cells were cultured in DMEM/F12 supplemented with 10% (v/v) FBS, 1% (v/v) penicillin-streptomycin, 1% Glutamax, 0.5ug/ml Amphotericin B and 0.26% sodium bicarbonate. To generate p53-/-clones by CRISPR-Cas9 mediated gene editing, RPE-1 cells were electroporated with Cas9:sgRNA ribonucleoparticles targeting the sequences 5’-AAATTTGCGTGTGGAG TATT-3’ and 5’-TCCACTCGGATAAGATGCTG-3’^52^ using the Neon Transfection system as described (Cirillo et al, 2024). After four days, cells were single sorted into 96-well plates containing medium supplemented with 10 µM nutlin-3a. After 14 days, surviving clones were expanded, and p53 status analysed by immunoblotting and sequencing of the TP53 locus as described^52^. CRISPR-Cas9 mediated gene editing of human RAP1 in p53-/-RPE-1 cells was carried out using phosphorothioated single-stranded DNA repair templates (ssODN) and selection for positive integrands by ouabain as described^53,54^. Briefly, RAP1 guide RNA 5’-GGCCCAGCCCGGCCAAGCGT-3’ was cloned into the BspI site of Addgene vector 86613. Repair templates for ATPA1 and RAP1 were synthesised by Integrated DNA Technologies (IDT) with the following sequences: ATP1A1: C*A*ATGTTACTGTGGATTGGAGCGATTCTTTGTTTCTTGGCTTATAGCATCAGA GCTGCTACAGAAGAGGAACCTCAAAACGATGACGTGAGTTCTGTAATTCAGCAT ATCGATTTGTAGTACACATCAGATATC*T*T. RAP1: C*A*TTCCTCGACTCTGTT CGTGAGGGACGACGGCAGCTCCATGTCCTTCTACGTGCGGCCCAGCCCGGCCGACGAGCGCCTCTCGACGCTCATCCTGCACGGCGGCGGCACGGTGTGCGAGGTGCAGGAGCCCGGGGCCGTGCTGCTGGCCCAGCCCGGGGAGGCGCTGGCCG AGGCCTCGGGTGATTTCATCTCCACG*C*A. where * denotes a phosphorthiolated base. To generate RAP1 KR/DE clones, 300,000 p53-/-RPE-1 cells were electroporated with the Neon Transfection System using a 10 µl tip and two pulses at 1350 V and 20 ms with 500 ng RAP1 gRNA/Addgene: 86613 plasmid, 2 pmol of ATP1A1 ssODN and 6 pmol RAP1 ssODN. To generate RAP1-/-clones, the procedure was repeated but omitting RAP1 ssODN. After 3-4 days, cells were expanded into 15 cm dished and treated with 0.25 µM ouabain, followed by isolation of single clones 7-12 days after drug selection. Genomic DNA was prepared using EZNA Tissue DNA kit according to the manufacturer’s instructions. *RAP1* was amplified by PCR using oligo sequences AGTGCTGCGCTTCGCGGC and CGCCTTCCGCTTGAGCTTCTG. Editing was analysed by restriction digestion with SalI and Sanger sequencing, and positive clones single cell sorted and expanded prior to freezing. For retro or lentiviral transduction, a total of 20 µg of plasmid DNA was transfected into Phoenix eco or 293T cells, respectively, using CaPO_4_ precipitation. The viral supernatant was filtered through a 0.45-μm filter, supplemented with 4 μg/ml polybrene, and used for the transduction of target cells. Lentiviral particles containing the sgRNA against mouse Rap1 (target: 5’-GCAGTCTAGGATGTACTGCG-3’) in lentiCRISPR v2 (Addgene plasmid # 52961, a gift from Feng Zhang) were introduced into target MEFs cells with three infections/day (6-12 h intervals) over 2 days, followed by 2 days in 2-4 µM Puromycin or 340 µM Hygromycin. The same approach was also used to target human Apollo in p53-/-RPE-1 cells with lentiviral particles containing lentiCRISPR v2 with the guide sequence 5’-CTGGTTCCAACGCAGCATGT-3’^23^, or non-targeting control sequence 5’-CGCCAAACGTGCCCTGACGG-3’. Cre was induced with three infections/day (6-12 h intervals) over two days with pMMP Hit & Run Cre retrovirus produced in Phoenix eco cells. Time-point 0 was set 12 hrs after the first Hit & Run Cre infection. For the Rap1-complementation assay, *Apollo^F/F^* MEFs were transduced with retroviral particle containing *Rap1*, *Rap1^KR/DE^*or *Rap1^R130E^* cloned in Plpc vector for a total of 4 infections at 6–12-h intervals, selected for 2-3 days in 2-4 µM Puromycin, transduced with the sgRNA against mouse Rap1 cloned in LcV2-Hygro (Addgene plasmid # 91977, a gift from Joshua Mendell), selected for 2 days in 340 µM Hygromycin and then transduced with pMMP Hit & Run Cre as previously described. For the Trf2-complementation assays, *Trf2*^F/F^*Rosa26Cre-ER*^T1^ MEFs were transduced with retroviral particle containing the Trf2 mutants, selected for 2 days in 2-4 µM Puromycin and then treated with 1 μM 4-OHT for 24 h. Time-point 0 was set at the time of 4-OHT addition.

### Generation of Trf2 mutant alleles

PCR was used to delete the Rap1 binding motif (RBM) or insert the S367A mutation in the MYC-tagged Trf2, Trf2^F120A^, Trf2^ΔiDDR^ and Trf2^F120A^ ^ΔiDDR^ alleles cloned in pLPC retroviral vector (Chen et al, 2008; Myler and Toia et al, 2023) using the following primers: Trf2^ΔRBM^-F: 5′-AATCTGGCATCCCCATCATCAC −3′ Trf2^1′RBM^-R: 5′-TCTG CTTGGAGGCTCTCTAAG −3′ Trf2^S367A^-F: 5′-GCGCCAGCCCACAAACACAAGAGA CC −3′ Trf2^S367A^-R: 5′-TGATGGGGATGCCAGATTAGCAAG-3′

### Fluorescence In Situ Hybridization (FISH)

FISH on mouse and human cells was performed as previously described^55^. Briefly, cells were treated treated with 0.2 µg/ml Colcemid (Biowest/Roche) for 2 hours before collection by trypsinization. Harvested cells were swollen in a hypotonic solution of 0.075M KCl at 37 °C for 20 minutes before fixation in methanol/acetic acid (3:1) overnight at 4 °C. Cells were dropped onto glass slides and allowed to age overnight. The slides were then dehydrated through an ethanol series of 70%, 95%, and 100% and allowed to air-dry. Telomere ends were hybridized with Cy3-OO-(TTAGGG)3 in hybridization solution (70% formamide, 1 mg/ml blocking reagent (1109617601, Roche), and 10 mM Tris-HCl pH 7.2) for 2 hours following an initial 5-10 min denaturation step at 80°C, washed twice with 70% formamide; 0.1% BSA; 10 mM Tris-HCl, pH 7.2 for 15 minutes each, and thrice in 0.08% Tween-20; 0.15 M NaCl; 0.1 M Tris-HCl, pH 7.2 or PBS for 5 minutes each. Chromosomal DNA was counterstained with the addition of DAPI (D1306, Invitrogen) to the second wash. Slides were left to air-dry and mounted in antifade reagent (Prolong Gold Antifade P36934, Fisher). coFISH analysis was performed as previously described^23^ with the following modifications: RPE-1 cells were treated with BrdU:BrdC for 14 h, slides were treated with UV for 10 minutes using a Blak ray model UV-21 365 nm handheld lamp at a distance of 8 cm and telomeres were detected with Cy3-OO-(TTAGGG)3 and FITC-OO-(CCCTAA)3.

### Immunoblotting

Cells were lysed in 2 x Laemmli buffer at 5×103 or 1×104 cells/μl and the lysate was denatured for 10 min at 95 °C before shearing with an insulin needle or sonication. Lysate equivalent to 1-2×105 cells was resolved using SDS/PAGE and transferred to a nitrocellulose membrane. Western blot was performed with 5% milk in PBS containing 0.1% (v/v) Tween-20 (PBS-T) with reagents described in the ‘antibodies used’ section. Signal was detected according to the manufacturer’s instructions using chemiluminescence western blotting detection reagents (Cytiva) using ChemiDoc (Bio-Rad) imaging systems.

### Chemical crosslinking mass spectrometry analysis

Complex assembly and crosslinking with DSSO was performed as described in the ‘Crosslinking’ section. After the crosslinking reaction, triethylammonium bicarbonate buffer (TEAB) was added to the sample at a final concentration of 100 mM. Proteins were reduced and alkylated with 5 mM tris-2-carboxyethyl phosphine (TCEP) and 10 mM iodoacetamide (IAA) simultaneously and digested overnight with trypsin at final concentration 50 ng/μL (Pierce). Sample was dried and peptides were fractionated with high-pH Reversed-Phase (RP) chromatography using the XBridge C18 column (1.0 × 100 mm, 3.5 μm, Waters) on an UltiMate 3000 HPLC system. Mobile phase A was 0.1% v/v ammonium hydroxide and mobile phase B was acetonitrile, 0.1% v/v ammonium hydroxide. The peptides were fractionated at 70 μL/min with the following gradient: 5 minutes at 5% B, up to 15% B in 3 min, for 32 min gradient to 40% B, gradient to 90% B in 5 min, isocratic for 5 minutes and re-equilibration to 5% B. Fractions were collected every 100 sec, SpeedVac dried and pooled into 12 samples for MS analysis. LC-MS analysis was performed on an UltiMate 3000 UHPLC system coupled with the Orbitrap Ascend Mass Spectrometer (Thermo). Each peptide fraction was reconstituted in 30 μL 0.1% formic acid and 15 μL were loaded to the Acclaim PepMap 100, 100 μm × 2 cm C18, 5 μm trapping column at 10 μL/min flow rate of 0.1% formic acid loading buffer. Peptides were then subjected to a gradient elution on the Acclaim PepMap (75 μm × 50 cm, 2 μm, 100 Å) C18 capillary column connected to the EASY-Spray source at 45 °C with an EASY-Spray emitter (Thermo, ES991). Mobile phase A was 0.1% formic acid and mobile phase B was 80% acetonitrile, 0.1% formic acid. The gradient separation method at flow rate 300 nL/min was as follows: for 80 min gradient from 5%-35% B, for 5 min up to 95% B, for 5 min isocratic at 95% B, re-equilibration to 5% B in 5 min, for 5 min isocratic at 5% B. Precursors between 380-1,400 m/z and charge states 3-8 were selected at 120,000 resolution in the top speed mode in 3 sec and were isolated for stepped HCD fragmentation (collision energies % = 21, 27, 34) with quadrupole isolation width 1.6 Th, Orbitrap detection with 30,000 resolution and 70 ms Maximum Injection Time. Targeted MS precursors were dynamically excluded from further isolation and activation for 45 seconds with 10 ppm mass tolerance. Identification of crosslinked peptides was performed in Proteome Discoverer 2.4 (Thermo) with the Xlinkx search engine in the MS2 mode for DSSO / +158.004 Da (K). Precursor and fragment mass tolerances were 10 ppm and 0.02 Da respectively with maximum 2 trypsin missed cleavages allowed. Carbamidomethyl at C was selected as static modification. Spectra were searched against a UniProt FASTA file containing Homo sapiens reviewed entries. Crosslinked peptides were filtered at FDR<0.01 using the Percolator node and target-decoy database search.

## Author contributions

M.E.D conceived the study, with genetic experiments devised in collaboration with F.L. Mouse embryonic fibroblast experiments were performed by C.S. with help from A.d.S. RPE-1 cell experiments were performed by M.E.D and P.E. with advice from J.M. Experiments in figures 2, 3 and 5 were performed by P.E with help from C.E.L.F. For figure 2b, S.G. designed the expression construct for shelterin, which was purified by O.I. Samples for figure 4 were prepared by P.E and C.E.L.F; O.I prepared and screened grids and P.E and C.E.L.F performed data processing. T.I.R performed cross-linking mass spectrometry for figure S5, on a sample prepared by P.E. Supervision was provided by J.S.C, S.G, F.L and M.E.D. The manuscript was written by M.E.D, with input from all authors.

## Acknowledgements

The authors are grateful to Peter Martin, Ronan Broderick, Alex Radzisheuskaya and Hannah Mischo for reagents and advice and to Gideon Coster and Titia de Lange for comments on the manuscript. M.E.D is funded by a Cancer Research UK Career Development Award (C68409/A28129). F.L. is supported by grants from Cancerfonden (21 1732 Pj), Vetenskapsrådet (2021-02788) and Knut and Alice Wallenberg Foundation. J.M. is funded by a Cancer Research UK Senior Cancer Research Fellowship (RCCSCF-Nov22/100001).

## Materials and Correspondence

Correspondence and requests for materials should be addressed to M.E.D and F.L.

## Supplementary figure legends

**Figure S1.** *RAP1 and APOLLO redundantly prevent cNHEJ at telomeres in mouse and human cells.* **a.** Immunoblot showing effective removal of Rap1 and persistence of TRF2 as indicated after Crispr-mediated Rap1 deletion and/or Cre-mediated deletion of Apollo in ApolloF/F Lig4+/+ or ApolloF/F Lig4-/-MEFs. Beta-actin is shown as loading control. **b.** Immunoblot showing expression of TRF2 in SV40LT-immortalized TRF2F/F RsCre-ERT1 MEFs transduced with an empty vector control (-) or mouse TRF2 and TRF2 F120A alleles deleted of the RAP1 binding motif (ΔRBM) and/or the DNA-damage response motif (ΔiDDR) 120 h after deletion of endogenous TRF2 with 4-OHT. Beta-actin shown as loading control. **c.** Representative FISH of metaphase spreads from cells as described in (b). Telomeres were detected with Cy3-(TTAGGG)3 (green). DNA was stained with DAPI (magenta). White and green arrows highlight chromatid-and chromosome type fusions respectively. **d.** Quantification of percentage telomeres involved in chromatid and chromosome fusions per metaphase after deletion of endogenous TRF2 in TRF2F/F RsCre-ERT1 MEFs expressing the indicated TRF2 mutants as described in (b). Data from 3 independent experiments, 10 metaphases per experiment (n = 30 total), with median. Statistics by ordinary One-way ANOVA. ‘ns’ not significant, ****P ≤ 0.0001. **e.** Sequencing of the RAP1 locus in p53-/-RAP1+/+ and p53-/-RAP1–/– RPE-1 cells. 19 bp deletion in knockout clones generates an altered reading frame that terminates in a stop codon after a total of 135 amino acids **f.** Immunoblot showing Apollo protein levels 120 h after transduction of p53-/-RAP1+/+ and p53-/-RAP1-/-RPE-1 cells with Cas9 and sgApollo or sgControl. **g.** chromatid-and chromosome-type telomere fusions quantified after deletion of Apollo as in (f) from three independent experiments, 10 metaphases from each, with median. Statistics as in (d) **h.** As in (g) but cells were treated with 2 µM DNA-PK inhibitor NU7741 for 48 prior to cell collection. From two independent experiments, 10 metaphases each, with median. Statistics as in (d) **i.** Representative coFISH images of RAP1 KO clone 2 120 h after transduction with Cas9 and sgApollo. Telomeres detected with Cy3-(TTAGGG)3 (red = leading) and FITC-(CCCTTA)3 (green = lagging). Arrows highlight leading:leading telomere fusions.

**Figure S2.** *Deletion constructs, purified proteins and data relating to figures 2 and 3.* **a.** Purified proteins as indicated, separated on a denaturing tris-glycine polyacrylamide gel and stained with Instant Blue. **b.** DNase I footprinting performed as in Fig. 2a with shelterin or TRF2/RAP1 at increasing concentrations (left panel) or with KU70/80, DNA-PKcs and TRF2/RAP1 each at 250 nM (right panel). Template concentation increased to 20 nM for the experiment on the right. **c.** DNase I footprinting performed as in Fig. 2a, except shelterin was added coincident with KU70/80 and DNA-PKcs. shelterin - POT1/TPP1 separated by SDS-PAGE and stained with Instant Blue. Cleavage products within the 10 bp footprint derive from a change in template sequence register compared with the standard template used in Figs. 2-4 **d.** Domain organisation of RAP1 and TRF2 mutants. TRF2 L330R (equivalent to L288R in the shorter TRF2 isoform) prevents binding to RAP1 (Chen et al, 2011). PreScission cleavage site highlighted by 3C. **e.** Purified proteins as indicated, separated on a denaturing tris-glycine polyacrylamide gel and stained with Instant Blue. **f.** Electrophoretic mobility shift assay of telomeric DNA bound by TRF2. **g.** DNaseI footprinting of DNA end-binding complexes. Proteins omitted as indicated. Telomeric DNA was used unless otherwise specified. **h.** Cleavage of Teb1 DBD from RAP1 using PreScission protease. Proteins were separated on a denaturing tris-glycine polyacrylamide gel and stained with Instant Blue.

**Figure S3.** *Cryo-EM data processing pipeline for the RAP1:DNA-PK complex on DNA.* Schematic showing the Cryospark classification and refinement steps used to obtain the RAP1:DNA-PK structure.

**Figure S4.** *DNA-PKcs conformation in the RAP1:DNA-PK complex.* DNA-PKcs density extracted from the structures indicated, showing the M-HEAT and N-HEAT domains adopting an ‘inactive’ conformation in the telomere end binding complex.

**Figure S5.** *Binding of the KU70 SAP and RAP1 myb domains to DNA.***a.** Density map and model of the SAP:DNA interaction extracted from the complete structure, showing K575, K595 and K596 coordinating the phosphate backbone **b.** Density map and model of the RAP1 myb:DNA interaction extracted from the complete structure, showing the recognition helix sitting in the major groove, and R133 acting as an N-terminal arm inserted into the neighbouring minor groove. Region protected from DNase I indicated **c.** Sequence alignment of the human RAP1 myb domain. Alignment adopts Clustal X colouring **d.** Comparison of the DNA-bound human RAP1 myb domain (blue) with homeotic protein antennapedia (purple - 1ahd) and HOX-B1 (pink - 1b72). Structures were aligned via the DNA. **e.** Table of intermolecular chemical crosslinks detected by XLMS. Data were thresholded with an XlinkX score ≥90 and crosslinks between KU and DNA-PKcs were excluded. **f.** Intermolecular chemical crosslinks detected by XLMS. Data were thresholded with an XlinkX score ≥90. **g.** Density map and model of the RAP1 loop C-terminal to the myb domain anchored to KU80.

**Figure S6.** *Binding of the RAP1 BRCT domain to KU70 vWA.* **a.** Sequence alignment of the RAP1 (left), KU70 (middle) and KU80 (right) regions that engage in the BRCT:KU interaction. Asterisks mark K39, R40 and R55 in RAP1, which are changed to aspartate or glutamate in the KR/DE mutant. Also, D496 and E499 in KU70 and D326 in KU80, which are changed to lysine or arginine in the KU DE/KR mutant. Alignment adopts Clustal X colouring **b.** Nano-scale differential scanning fluorimetry analysis of the RAP1 BRCT or KU mutants as indicated, showing no significant effect of the point mutations on protein folding. **c.** Protein cross-linking analysis of RAP1 and KU in the presence of DNA. Proteins were mixed with crosslinker and the products of the reaction were separated on a denaturing tris-acetate polyacrylamide gel and analysed by silver staining or immunoblotting as indicated. RAP1 KR/DE (KR) contains K39D, R40E and R55E mutations. KU DE/KR (DE) contains KU70 D496K, E499R and KU80 D327K mutations.

**Figure S7.** *Supplementary DNA-PK binding assays related to* figure 5. **a.** KU pulldown experiment without TRF2 or RAP1, executed as described in Figure 5a. KU was immunoprecipitated from reactions containing KU70/80, DNA-PKcs, XRCC4/LIG4 and DNA template as indicated, separated on a denaturing polyacrylamide gel and immunoblotted as indicated. KU DE/KR contains the mutations KU70 D496K, E499R and KU80 D327K **b.** As in (a), but with reactions containing 5 nM TRF2/RAP1 complex and increasing concentrations of XRCC4/LIG4 as indicated. **c.** as in (b), but with the XRCC4/LIG4 concentrations indicated incubated with KU70/80, DNA-PKcs and DNA template for 5 minutes prior to the addition of TRF2/RAP1. **d.** As in (b), but with 30 nM XRCC4/LIG4, and the RAP1 proteins indicated.

**Figure S8.** *Mutation analysis of RAP1 in mouse and human cells* **a.** Immunoblot showing over-expression of mouse RAP1, RAP1 KR/DE and RAP1 R130E after Crispr-and Cre-mediated deletion of Rap1 and Apollo respectively in ApolloF/F Lig4+/+ MEFs. Beta-actin is shown as loading control. **b.** Representative FISH of metaphase spreads of ApolloF/F Lig4+/+ MEFs expressing the indicated RAP1 mutants or a control empty vector (EV) 96-120 h after deletion of endogenous Rap1 with sgRNA and 120 h after deletion of APOLLO with Hit & Run Cre. Telomeres were detected with Cy3-(TTAGGG)3 (green). DNA was stained with DAPI (magenta). White and green arrows highlight chromatid-and chromosome type fusions respectively. **c.** Quantification of the percentage of telomeres involved in chromatid fusions per metaphases after expression of Rap1 or the empty vector control (-) and removal of endogenous Rap1 and Apollo as described in (b). Data from 3 independent experiments, 10 metaphases per experiment (n = 30 total), with median. Statistics by ordinary One-way ANOVA. **d.** Sequence analysis of the human RAP1 locus in RAP1 WT and RAP1 KR/DE p53-/-RPE-1 cells. Targeted mutations in KR/DE are marked with an asterisk, with altered amino acids highlighted in red **e.** Immunoblot showing Apollo levels after Crispr-mediated deletion in p53-/-RAP1 WT and p53-/-RAP1 KR/DE RPE-1 cells. Asterisk marks a non-specific band detected by the anti-Apollo antibody. Tubulin as loading control **f.** Representative FISH of metaphase spreads of RAP1 WT and RAP1 KR/DE p53–/– RPE-1 cells 120 h after transduction with Cas9 and sgApollo or sgControl as indicated. Telomeres detected with Cy3-(TTAGGG)3 (green). DNA stained with DAPI (magenta). White and green arrows highlight chromatid-and chromosome type fusions respectively. **g.** Quantification of the percentage of telomeres involved in chromatid and chromosome fusions per metaphases after removal of Apollo in RAP1 WT and RAP1 KR/DE p53–/– RPE-1 cells as indicated. Data from 3 independent experiments, 10 metaphases per experiment (n = 30 total), with median. Statistics by ordinary One-way ANOVA.

**Figure S9.** *Conservation analysis of the RAP1 BRCT:KU interaction*. Amino acid sequences of RAP1 from the metazoan species indicated were aligned using the Muscle algorithm. The basic patch composed of K39, R40, R41 and R55 in the BRCT domain is shown with basic residues displayed in red. Corresponding structure predictions for the RAP1:KU complex made using Alphafold 3 are shown for the species indicated, revealing that the BRCT:KU interaction is likely to occur widely across metazoa.

**Supplementary Video 1.** Video showing the cryo-EM composite map of the RAP1:DNA-PK complex with annotation.

