## Supplementary data for "Chromosome end protection by RAP1-mediated inhibition of DNA-PK"

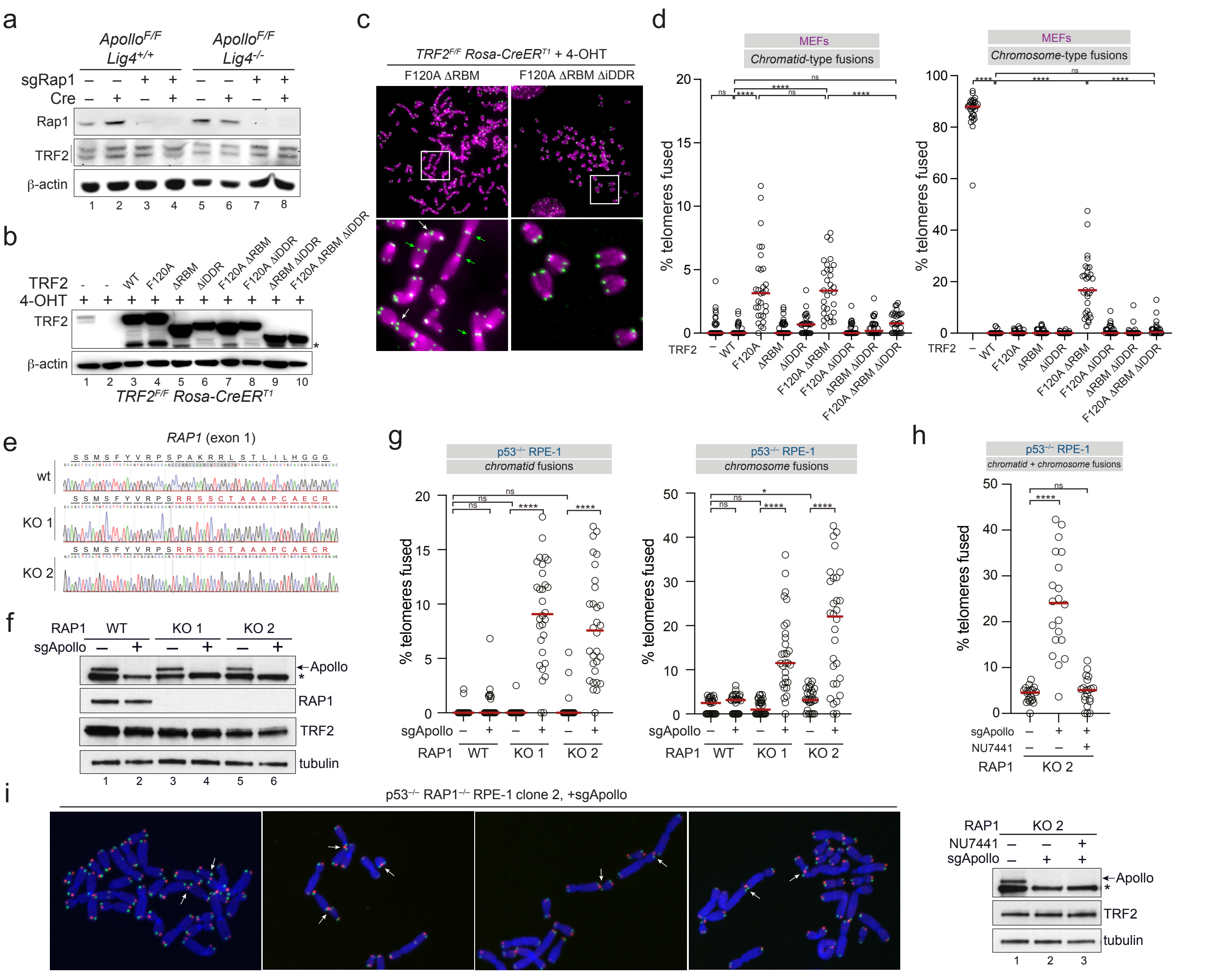

***Figure S1. RAP1 and APOLLO redundantly prevent cNHEJ at telomeres in mouse and human cells.*** **a.** Immunoblot showing effective removal of Rap1 and persistence of TRF2 as indicated after Crispr-mediated Rap1 deletion and/or Cre-mediated deletion of Apollo in ApolloF/F Lig4+/+ or ApolloF/F Lig4-/- MEFs. Beta-actin is shown as loading control. **b.** Immunoblot showing expression of TRF2 in SV40LT-immortalized TRF2F/F RsCre-ERT1 MEFs transduced with an empty vector control (-) or mouse TRF2 and TRF2 F120A alleles deleted of the RAP1 binding motif (ΔRBM) and/or the DNA-damage response motif (ΔiDDR) 120 h after deletion of endogenous TRF2 with 4-OHT. Beta-actin shown as loading control. **c.** Representative FISH of metaphase spreads from cells as described in (b). Telomeres were detected with Cy3-(TTAGGG)3 (green). DNA was stained with DAPI (magenta). White and green arrows highlight chromatid- and chromosome type fusions respectively. **d.** Quantification of percentage telomeres involved in chromatid and chromosome fusions per metaphase after deletion of endogenous TRF2 in TRF2F/F RsCre-ERT1 MEFs expressing the indicated TRF2 mutants as described in (b). Data from 3 independent experiments, 10 metaphases per experiment (n = 30 total), with median. Statistics by ordinary One-way ANOVA. ‘ns’ not significant, ****P ≤ 0.0001. **e.** Sequencing of the *RAP1* locus in p53-/- RAP1+/+ and p53-/- RAP1–/– RPE-1 cells. 19 bp deletion in knockout clones generates an altered reading frame that terminates in a stop codon after a total of 135 amino acids **f.** Immunoblot showing Apollo protein levels 120 h after transduction of p53-/- RAP1+/+ and p53-/- RAP1-/- RPE-1 cells with Cas9 and sgApollo or sgControl. **g.** chromatid- and chromosome-type telomere fusions quantified after deletion of Apollo as in (f) from three independent experiments, 10 metaphases from each, with median. Statistics as in (d) **h.** As in (g) but cells were treated with 2 μM DNA-PK inhibitor NU7741 for 48 prior to cell collection. From two independent experiments, 10 metaphases each, with median. Statistics as in (d) **i.** Representative coFISH images of RAP1 KO clone 2 120 h after transduction with Cas9 and sgApollo. Telomeres detected with Cy3-(TTAGGG)3 (red = leading) and FITC-(CCCTTA)3 (green = lagging). Arrows highlight leading:leading telomere fusions.

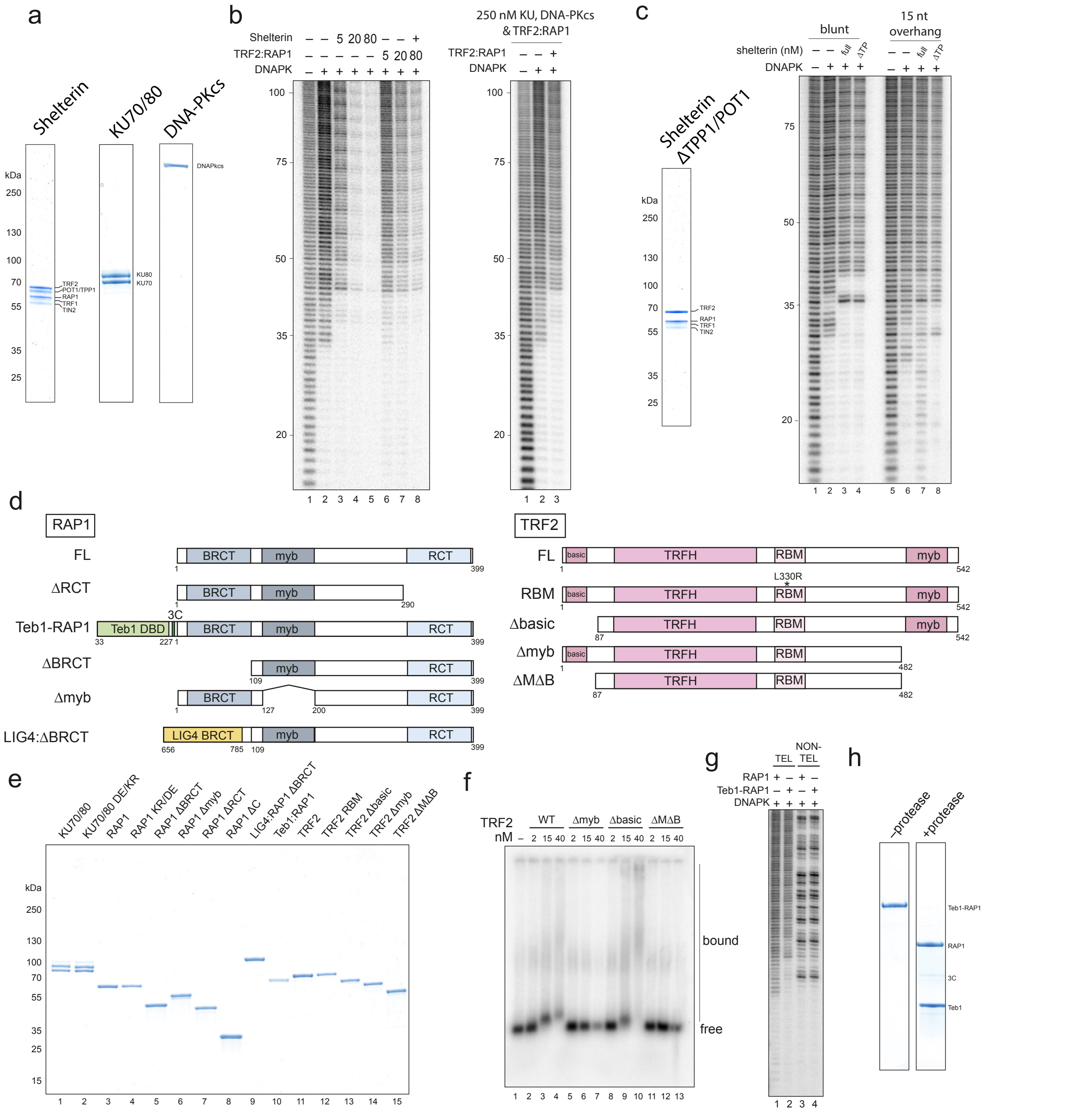

***Figure S2. Deletion constructs, purified proteins and data relating to figures 2 and 3.*** **a.** Purified proteins as indicated, separated on a denaturing tris-glycine polyacrylamide gel and stained with Instant Blue. **b.** DNase I footprinting performed as in Fig. 2a with shelterin or TRF2/RAP1 at increasing concentrations (left panel) or with KU70/80, DNA-PKcs and TRF2/RAP1 each at 250 nM (right panel). Template concentation increased to 20 nM for the experiment on the right. **c.** DNase I footprinting performed as in Fig. 2a, except shelterin was added coincident with KU70/80 and DNA-PKcs. shelterin -POT1/TPP1 separated by SDS-PAGE and stained with Instant Blue. Cleavage products within the 10 bp footprint derive from a change in template sequence register compared with the standard template used in Figs. 2-4 **d.** Domain organisation of RAP1 and TRF2 mutants. TRF2 L330R (equivalent to L288R in the shorter TRF2 isoform) prevents binding to RAP1 (Chen et al, 2011). PreScission cleavage site highlighted by 3C. **e.** Purified proteins as indicated, separated on a denaturing tris-glycine polyacrylamide gel and stained with Instant Blue. **f.** Electrophoretic mobility shift assay of telomeric DNA bound by TRF2. **g.** DNaseI footprinting of DNA end-binding complexes. Proteins omitted as indicated. Telomeric DNA was used unless otherwise specified. **h.** Cleavage of Teb1 DBD from RAP1 using PreScission protease. Proteins were separated on a denaturing tris-glycine polyacrylamide gel and stained with Instant Blue.

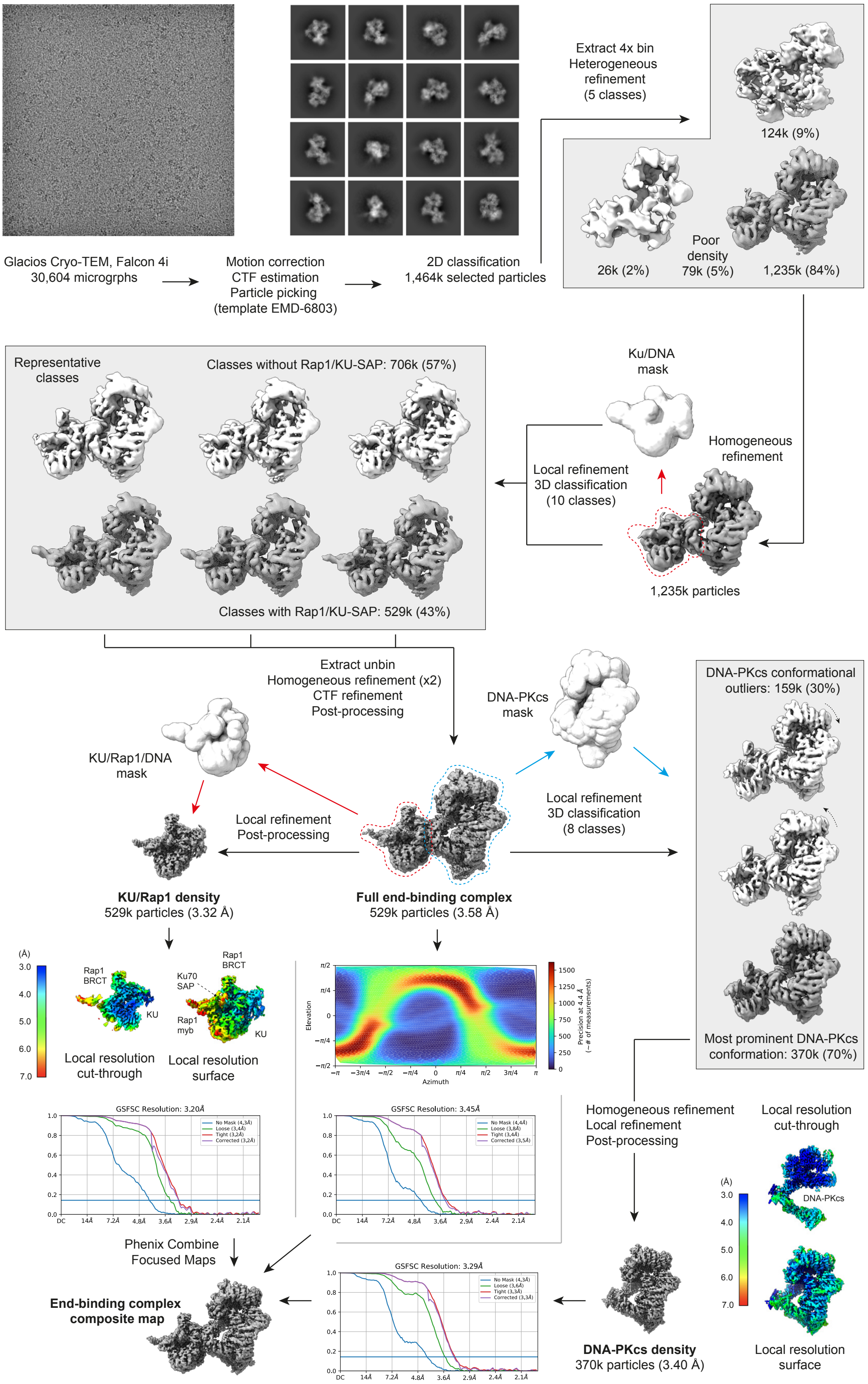


***Figure S3. Cryo-EM data processing pipeline for the RAP1:DNA-PK complex on DNA.*** Schematic showing the Cryospark classification and refinement steps used to obtain the RAP1:DNA-PK structure.

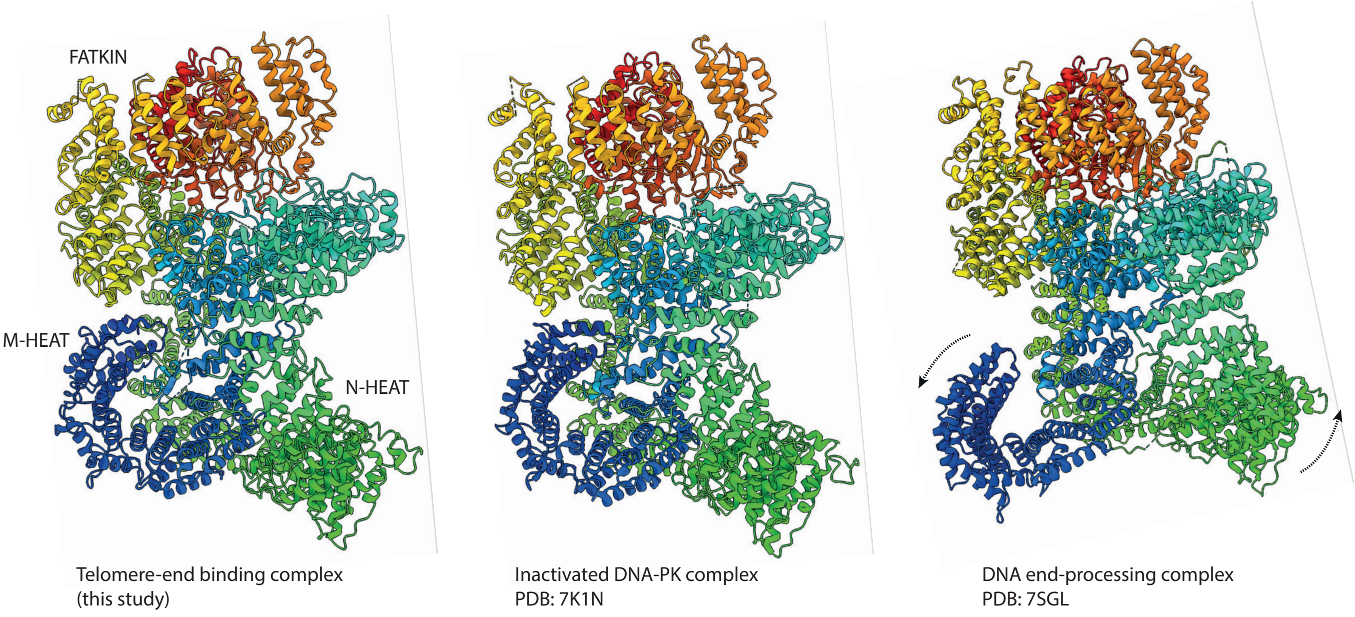


***Figure S4. DNA-PKcs conformation in the RAP1:DNA-PK complex.*** DNA-PKcs density extracted from the structures indicated, showing the M-HEAT and N-HEAT domains adopting an 'inactive' conformation in the telomere end binding complex.

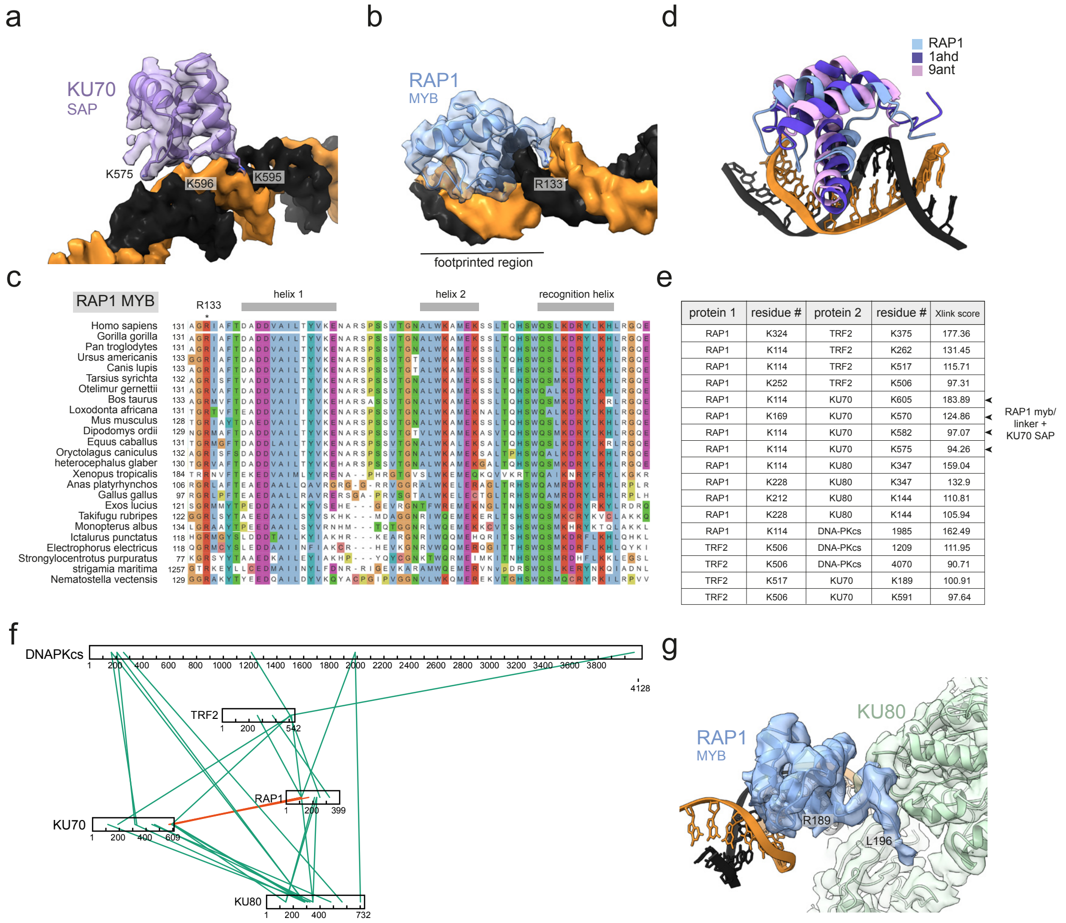


| protein 1 | residue # | protein 2 | residue # | Xlink score |
|---|---|---|---|---|
| RAP1 | K324 | TRF2 | K375 | 177.36 |
| RAP1 | K114 | TRF2 | K262 | 131.45 |
| RAP1 | K114 | TRF2 | K517 | 115.71 |
| RAP1 | K252 | TRF2 | K506 | 97.31 |
| RAP1 | K114 | KU70 | K605 | 183.89 |
| RAP1 | K169 | KU70 | K570 | 124.86 |
| RAP1 | K114 | KU70 | K582 | 97.07 |
| RAP1 | K114 | KU70 | K575 | 94.26 |
| RAP1 | K114 | KU80 | K347 | 159.04 |
| RAP1 | K228 | KU80 | K347 | 132.9 |
| RAP1 | K212 | KU80 | K144 | 110.81 |
| RAP1 | K228 | KU80 | K144 | 105.94 |
| RAP1 | K114 | DNA-PKcs | 1985 | 162.49 |
| TRF2 | K506 | DNA-PKcs | 1209 | 111.95 |
| TRF2 | K506 | DNA-PKcs | 4070 | 90.71 |
| TRF2 | K517 | KU70 | K189 | 100.91 |
| TRF2 | K506 | KU70 | K591 | 97.64 |

***Figure S5. Binding of the KU70 SAP and RAP1 myb domains to DNA*** **a.** Density map and model of the SAP:DNA interaction extracted from the complete structure, showing K575, K595 and K596 coordinating the phosphate backbone **b.** Density map and model of the RAP1 myb:DNA interaction extracted from the complete structure, showing the recognition helix sitting in the major groove, and R133 acting as an N-terminal arm inserted into the neighbouring minor groove. Region protected from DNase I indicated **c.** Sequence alignment of the human RAP1 myb domain. Alignment adopts Clustal X colouring **d.** Comparison of the DNA-bound human RAP1 myb domain with other homeodomains as indicated. Structures were aligned via the DNA. **e.** Table of intermolecular chemical crosslinks detected by XLMS. Data were thresholded with an XlinkX score ≥90 and crosslinks between KU and DNA-PKcs were excluded. **f.** Intermolecular chemical crosslinks detected by XLMS. Data were thresholded with an XlinkX score ≥90. **g.** Density map and model of the RAP1 loop C-terminal to the myb domain anchored to the KU80 vWA domain

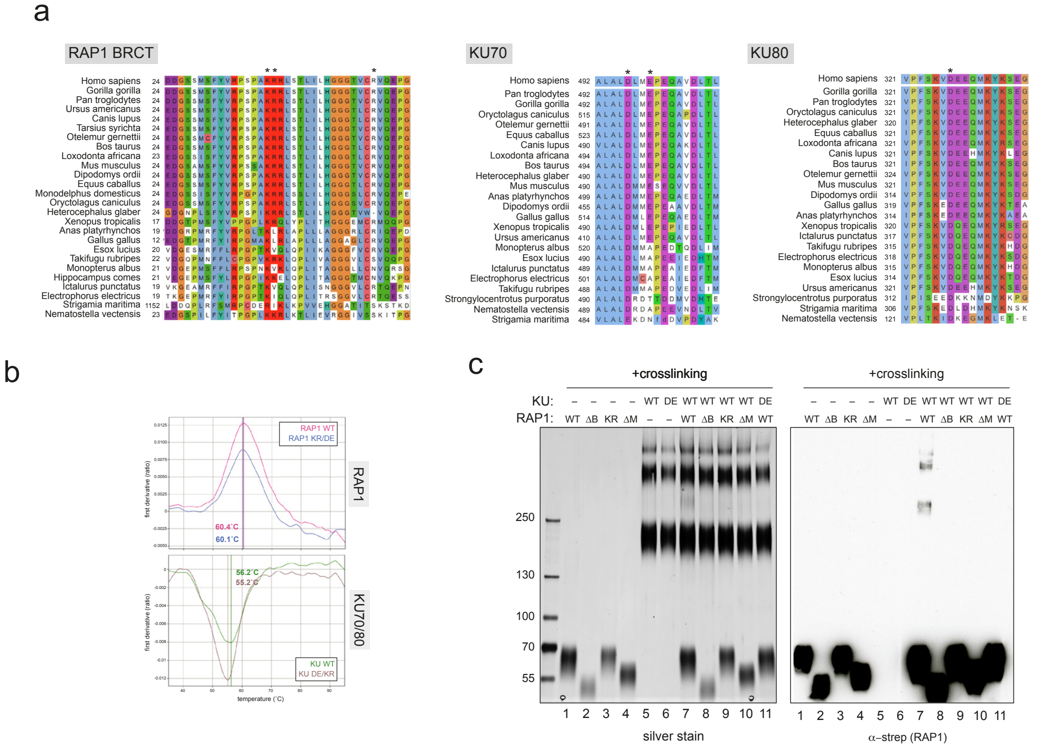


***Figure S6. Binding of the RAP1 BRCT domain to KU70 vWA.*** **a.** Sequence alignment of the RAP1 (left), KU70 (middle) and KU80 (right) regions that engage in the BRCT:KU interaction. Asterisks mark K39, R40 and R55 in RAP1, which are changed to aspartate or glutamate in the KR/DE mutant. Also, D496 and E499 in KU70 and D326 in KU80, which are changed to lysine or arginine in the KU DE/KR mutant. Alignment adopts Clustal X colouring **b.** Nano-scale differential scanning fluorimetry analysis of the RAP1 BRCT or KU mutants as indicated, showing no significant effect of the point mutations on protein folding. **c.** Protein cross-linking analysis of RAP1 and KU in the presence of DNA. Proteins were mixed with crosslinker and the products of the reaction were separated on a denaturing tris-acetate polyacrylamide gel and analysed by silver staining or immunoblotting as indicated. RAP1 KR/DE (KR) contains K39D, R40E and R55E mutations. KU DE/KR (DE) contains KU70 D496K, E499R and KU80 D327K mutations.

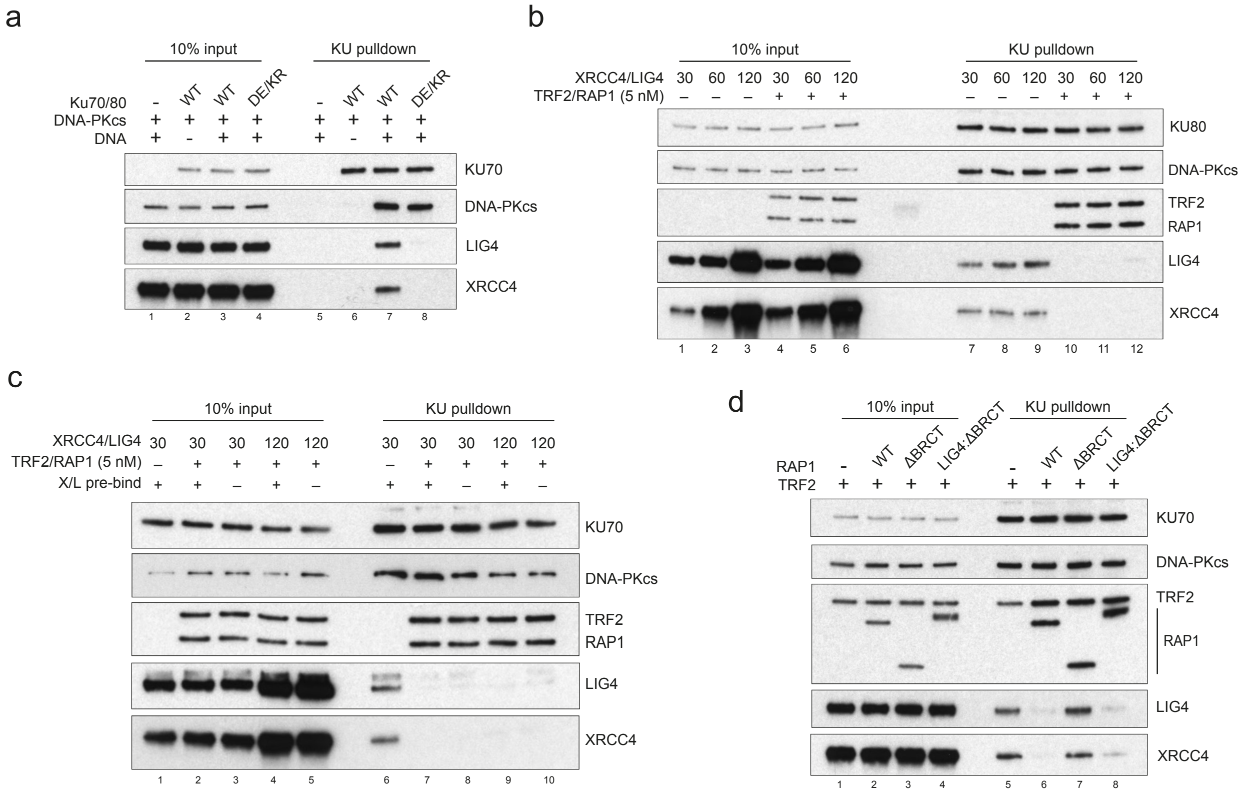


***Figure S7. Supplementary DNA-PK binding assays related to figure 5.*** **a.** KU pulldown experiment without TRF2 or RAP1, executed as described in Figure 5a. KU was immunoprecipitated from reactions containing KU70/80, DNA-PKcs, XRCC4/LIG4 and DNA template as indicated, separated on a denaturing polyacrylamide gel and immunoblotted as indicated. KU DE/KR contains the mutations KU70 D496K, E499R and KU80 D327K **b.** As in a., but with reactions containing 5 nM TRF2/RAP1 complex and increasing concentrations of XRCC4/LIG4 as indicated. **c.** as in b., but with the XRCC4/LIG4 concentrations indicated incubated with KU70/80, DNA-PKcs and DNA template for 5 minutes prior to the addition of TRF2/RAP1. **d.** As in b., but with 30 nM XRCC4/LIG4, and the RAP1 proteins indicated.

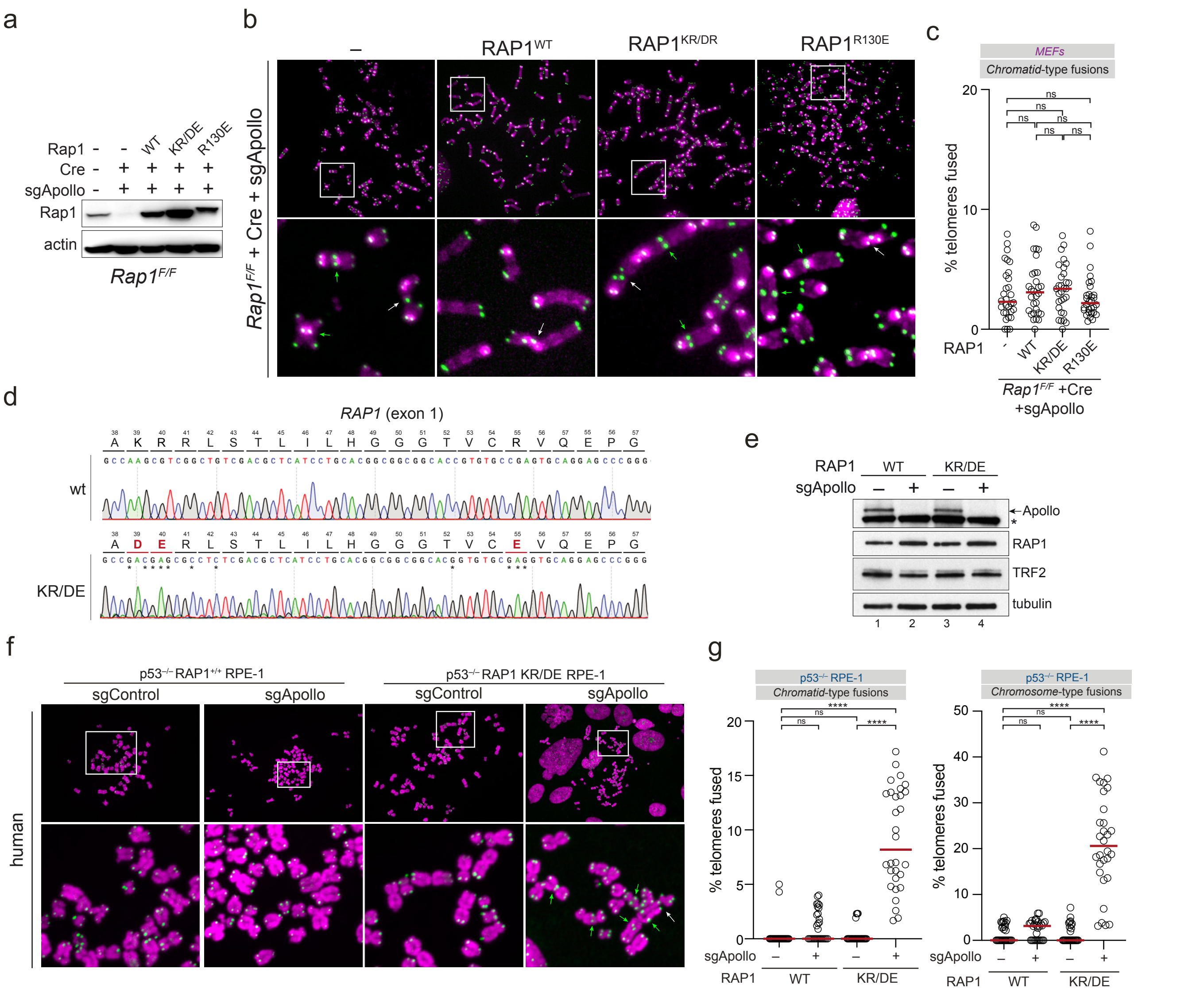


***Figure S8. Mutation analysis of RAP1 in mouse and human cells*** **a.** Immunoblot showing over-expression of mouse RAP1, RAP1 KR/DE and RAP1 R130E after Crispr- and Cre-mediated deletion of Rap1 and Apollo respectively in ApolloF/F Lig4+/+ MEFs. Beta-actin is shown as loading control. **b.** Representative FISH of metaphase spreads of ApolloF/F Lig4+/+ MEFs expressing the indicated RAP1 mutants or a control empty vector (EV) 96-120 h after deletion of endogenous Rap1 with sgRNA and 120 h after deletion of APOLLO with Hit & Run Cre. Telomeres were detected with Cy3-(TTAGGG)3 (green). DNA was stained with DAPI (magenta). White and green arrows highlight chromatid- and chromosome type fusions respectively. **c.** Quantification of the percentage of telomeres involved in chromatid fusions per metaphases after expression of Rap1 or the empty vector control (-) and removal of endogenous Rap1 and Apollo as described in b. Data from 3 independent experiments, 10 metaphases per experiment (n = 30 total), with median. Statistics by ordinary One-way ANOVA. **d.** Sequence analysis of the human *RAP1* locus in RAP1 WT and RAP1 KR/DE p53-/- RPE-1 cells. Targetted mutations in KR/DE are marked with an asterisk, with altered amino acids highlighted in red **e.** Immunoblot showing Apollo levels after Crispr-mediated deletion in p53-/- RAP1 WT and p53-/- RAP1 KR/DE RPE-1 cells. Asterisk marks a non-specific band detected by the anti-Apollo antibody. Tubulin as loading control **f.** Representative FISH of metaphase spreads of RAP1 WT and RAP1 KR/DE p53–/– RPE-1 cells 120 h after transduction with Cas9 and sgApollo or sgControl as indicated. Telomeres detected with Cy3-(TTAGGG)3 (green). DNA stained with DAPI (magenta). White and green arrows highlight chromatid- and chromosome type fusions respectively. **g.** Quantification of the percentage of telomeres involved in chromatid and chromosome fusions per metaphases after removal of Apollo in RAP1 WT and RAP1 KR/DE p53–/– RPE-1 cells as indicated. Data from 3 independent experiments, 10 metaphases per experiment (n = 30 total), with median. Statistics by ordinary One-way ANOVA.

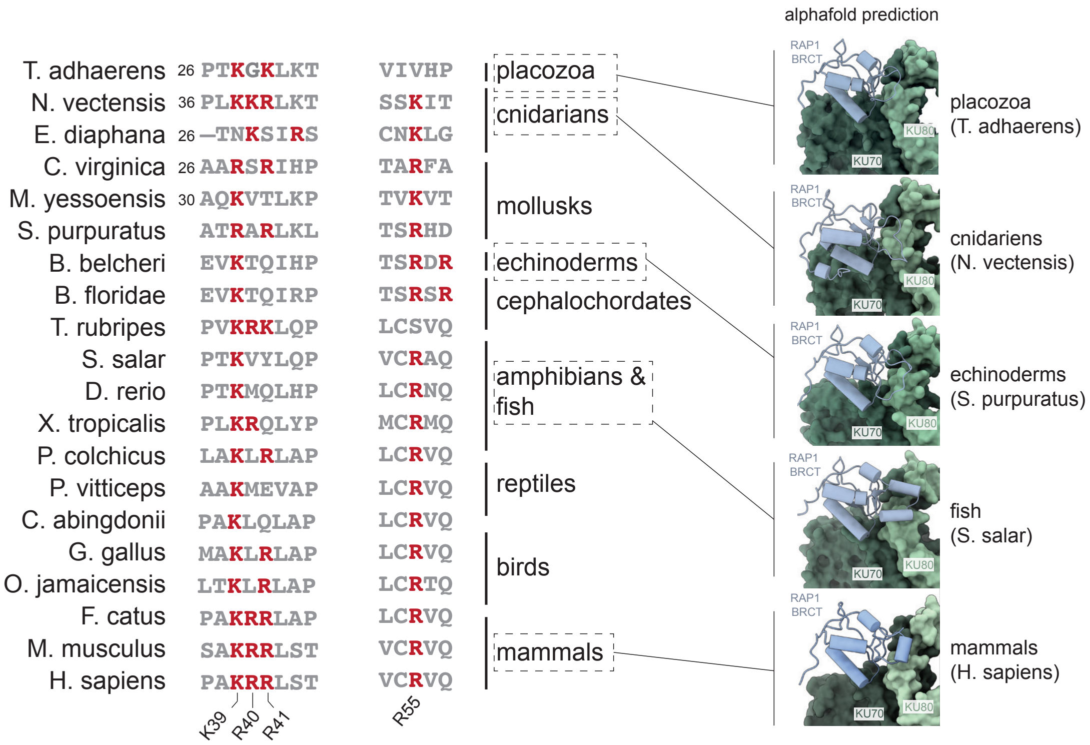


***Figure S9. Conservation analysis of the RAP1 BRCT:KU interaction*** **a.** Amino acid sequences of RAP1 from the metazoan species indicated were aligned using the Muscle algorithm. The basic patch composed of K39, R40, R41 and R55 in the BRCT domain is shown with basic residues displayed in red. Corresponding structure predictions for the RAP1:KU complex made using Alphafold 3 are shown for the species indicated, revealing that the BRCT:KU interaction is likely to occur widely across metazoa.
